# Axially decoupled photo-stimulation and two photon readout (*ADePT*) for mapping functional connectivity of neural circuits

**DOI:** 10.1101/2025.02.24.639992

**Authors:** Matthew Koh, Francesca Anselmi, Sanjeev K. Kaushalya, Diego E. Hernandez, Walter Germán Bast, Pablo S. Villar, Honggoo Chae, Martin B. Davis, Sadhu Sai Teja, Zhe Qu, Viviana Gradinaru, Priyanka Gupta, Arkarup Banerjee, Dinu F. Albeanu

**Author notes:** equal contribution (Matthew Koh, Francesca Anselmi, Sanjeev K. Kaushalya).

## Abstract

All-optical *in vivo* physiology enables three-dimensional interrogation of neural circuits function. Here, we introduce *ADePT* (Axially-Decoupled Photo-stimulation and Two-photon readout), a strategy that combines flexible optogenetic stimulation with neural activity imaging. *ADePT* achieves axially contained widefield patterned stimulation by coupling a digital micro-mirror device illuminated by a solid-state laser to a motorized holographic diffuser, while in parallel performing multiphoton imaging across different z-planes. As a proof of principle, we apply *ADePT* to map excitatory and inhibitory functional connectivity in the mouse early olfactory system. By controlling activity in individual glomeruli on the olfactory bulb surface and recording responses from output mitral and tufted cells in deeper layers, we identify cohorts of sister cells driven by the same parent glomerulus and analyze their distinct connectivity patterns. We further report specificity in the inhibitory circuitry, as we find that GABAergic/dopaminergic (DAT+) interneurons associated with different glomeruli selectively suppress baseline and odor-evoked activity in distinct mitral and tufted cell populations. Together, these results establish *ADePT* as a platform for high-throughput functional connectivity mapping in optically accessible brain regions.

## Introduction

Manipulating select neural circuit elements is critical for understanding the circuit mechanisms underlying brain computation, especially when coupled with activity monitoring and behavioral analysis. In recent years, optogenetic control of specific cell-types has been widely adopted as a method of choice^1–5^, and various combinations of optical imaging and spatial light patterning techniques^6–19^ have achieved concurrent stimulation and activity recording, typically in the same optical z-plane. This is primarily constrained by the microscope anatomy, as the photo-stimulation and imaging light paths converge onto the sample through a shared microscope objective which focuses both the stimulation and imaging light patterns in its fixed focal plane. Neuronal circuits are however distributed in 3-D and investigating the functional connectivity of such neural circuits requires flexible control and readout of neural activity in independent optical planes (e.g. stimulating layer 1 dendrites and monitoring neuronal activity in layers 5, **Fig. 1a**).

**Figure 1:**
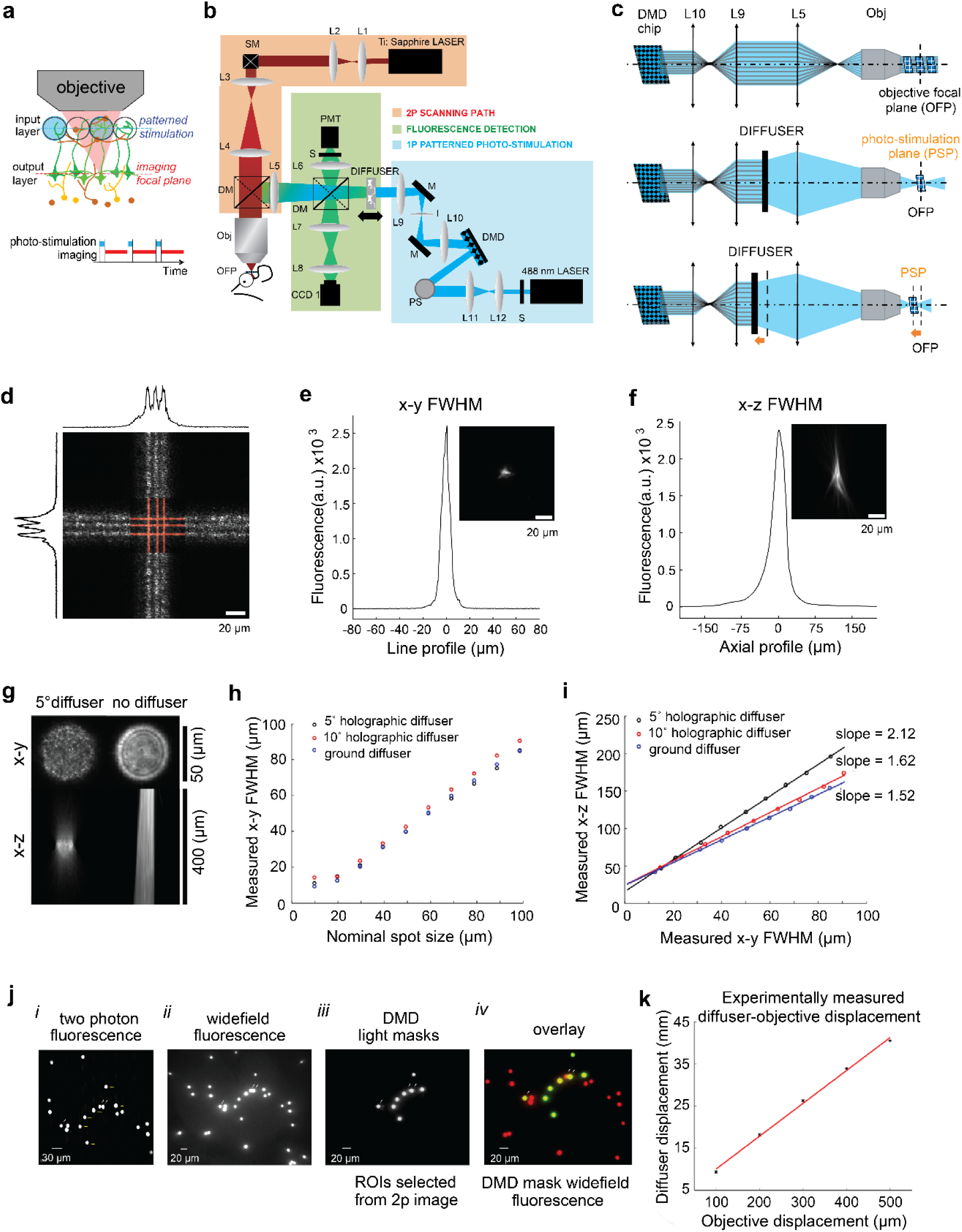
Axially decoupled patterned photo-stimulation and multiphoton imaging. **(a)** (Top) Cartoon schematics of functional connectivity mapping of a neural circuit (e.g. olfactory bulb); photo-stimulation (2-D light mask from a digital micro-mirror device illuminated with a solid-state CW laser) and two photon imaging are confined to different optical z-planes that can be flexibly and independently adjusted by translating the diffuser and respectively the primary objective; (Bottom) Alternating (strobing) between photo-stimulation and imaging periods. Each red bar represents a single frame of multiphoton imaging. Photo-stimulation and imaging periods are interleaved. (**b)** Microscope schematics. DM, dichroic mirrors. DMD, digital micro-mirror device. I, iris diaphragm. L1-L12, lenses. O, primary objective. PMT, photomultiplier tube. PS, periscope. S, shutter. SM, scan mirrors. **(c)** (Top) Illustration of using a movable diffuser to decouple the patterned photo-stimulation and multiphoton imaging planes. The diffuser is imaged into the sample, in a 4f lens configuration; translating the diffuser along the optical path causes the corresponding projection plane to shift axially. OFP, objective focal plane. PSP, photo-stimulation plane. **(d)** Example projected grid pattern (red overlay) and ensuing fluorescence on a fluorescently coated slide; line width is 1 pixel (2.4 µm); distance between lines is 2 and 3 pixels respectively; marginal distributions represent average cross-section line profiles. **(e)** x-y FWHM (6.2 ± 1.6 µm) of a 4.8 µm nominal diameter spot. (**f**) x-z FWHM (34.0 ± 4.9 µm) of a 4.8 µm nominal diameter spot. (**g)** Axial confinement of light intensity of a 2-D projection pattern using a holographic diffuser conjugated to the focal plane of the principal objective (Left) vs. same optical configuration without a diffuser and the resultant light pencil (Right); spot diameter = 50 µm. **(h)** Measured spot x-y FWHM versus expected nominal spot size calculated by multiplying the pattern size in pixels by the expected size of a pixel given the DMD chip and the optical magnification of the system. **(i)** Measured axial (z) FWHM versus lateral (x-y) FWHM of the projected spots from **(h)** for three different types of diffusers; black circles = 5° holographic diffuser; red circles = 10° holographic diffuser; blue circles = grit ground diffuser. **(j)** DMD chip-to-CCD camera-to-2p microscope registration. (i) two photon micrograph of 10 µm fluorescence microbeads; arrows mark two microbeads, part of the larger DMD-modulated projection target pattern (8 microbeads), which were taken as fiduciary points; (ii) widefield fluorescence image (full field illumination) of a larger field of view including the target microbeads; (iii) ROIs selected from the 2p image were used to generate DMD-chip light masks; these were further projected at the primary objective focal plane and imaged using the primary CCD camera (CCD 1); (iv) overlay of DMD-generated photo-stimulation masks and widefield fluorescence image of 10 µm microbeads from (ii); note that fluorescence is restricted to the microbeads targeted by the DMD photo-stimulation masks with minimal spillover to adjacent (off-target) microbeads (see fiduciary marks). **(k)** Measured calibration curve of diffuser translation required to shift photo-stimulation plane by a given amount with respect to the focal plane of the objective. Each point represents Mean ± SEM across ∼6 months, showing the robustness of the setup and linear relationship between objective and compensatory diffuser displacements for the range of decoupling between the photo-stimulation and imaging planes sampled (500 µm).

Several strategies have been developed over the past two decades to target multiple optical z-planes simultaneously, or in quick succession. These range from dual scanning head implementations^19^ to using tunable focusing elements (e.g. electrically-tunable lenses, ETL) whose focal length can be dynamically controlled^20–22^, acousto-optic deflector (AOD)-based arbitrary access imaging systems^14,23–28^ and digital holography^8,9,13,27,29–33^. These methods have varying trade-offs, and to date, monitoring and photo-activating neurons in three dimensions remains challenging; existing approaches are either cost-prohibitive, limited in field of stimulation, or introduce optical aberrations that compromise imaging quality. Here we introduce *ADePT* (**A**xially **De**coupled **P**hoto-stimulation and **T**wo-Photon readout), an easily implementable, cost-effective and high-throughput technique that combines one-photon widefield patterned illumination with multiphoton imaging in axially decoupled planes (**Fig. 1a-c**). This approach enables axially-restricted photo-stimulation with arbitrarily shaped light patterns across an extended region on the brain surface, within 100 µm from pia, while imaging neuronal activity in deeper optically accessible z-planes of choice. The DMD-based photo-stimulation approach is particularly well-suited for understanding information flow in areas where sensory stimuli evoke structured patterns of activity extending over several 10s of micrometers. We use a digital micro-mirror device (DMD)^17,34–47^ to impose customizable 2-D spatial patterns onto a Gaussian laser beam that are subsequently optically relayed onto the sample. Building on work from other groups^38,48–54^, we constrain light patterns using diffuser elements. To flexibly target photo-stimulation in different z-planes independent of the multiphoton imaging plane, we translate a diffuser along the optical path using a motorized stage. Akin to remote focusing imaging approaches^16,20,55–60^, translating the diffuser displaces the optical z-plane within the sample where the stimulation pattern is in focus independent of the objective’s focal plane (**Fig. 1c**). *ADePT* enables patterned photo-stimulation across a large field of stimulation, and can be calibrated across hundreds of micrometers of axial displacement between the stimulation and imaging planes. These parameters are readily adjustable to suit different operating configurations and biological applications.

Using *ADePT*, in proof-of-principle experiments, we investigate the logic of circuit connectivity by leveraging the 3-D spatial layout of the mouse olfactory bulb (OB)^61,62^. In the OB, the input nodes called glomeruli (clusters of neuropil sorted by odorant receptor identity) and the principal output neurons (mitral and tufted cells) are separated by approximately two hundred microns in depth. We photo-stimulated either the olfactory sensory neuron inputs to glomeruli or local inhibitory interneurons while monitoring responses of deeper mitral/tufted cells (MT cells). We find that *sister* OB output neurons^34,46,46,63–74^ – those that receive primary input from the *same* ‘parent’ glomerulus – differ in their lateral inhibitory connectivity with other glomeruli with the OB network, potentially explaining their non-redundant responses to odors^34,63^. Further, local GABAergic/dopaminergic interneurons^40,41,75–83^ associated with different glomeruli provide dense, but selective inhibition that sparsens and decorrelates the odor responses of OB output neurons. In summary, *ADePT* is readily-deployable for mapping excitatory and inhibitory functional connectivity in optically accessible brain regions when target structures are superficial and resolution requirements are supracellular.

## Results

### Design principle of *ADePT*

*ADePT* couples a custom laser scanning two photon microscope (Methods) with a modular add-on for axially-contained widefield (one-photon) patterned photo-stimulation (**Fig. 1b**). Patterned photo-stimulation is achieved using a digital micro-mirror device (VX4100 DMD, 0.7″ XGA VIS 1024 x 768 pixels, 13.68 µm micro-mirror pitch, Vialux) illuminated by a solid-state CW laser (Sapphire, 488 nm, 200 mW, Coherent). The photo-stimulation and two-photon excitation paths are combined using a dichroic mirror (680dcxr, 50 mm diameter, Chroma) placed above the objective (20X, 1.0 NA, Olympus). A diffuser mounted on a motorized translation stage enables flexible control of the axial location of the photo-stimulation plane in the sample (**Figs. 1a-c**). Without the diffuser, the photo-stimulation 2-D pattern is in focus throughout the optical path as pencils of coherent light (infinite focus, **Fig. 1c**, *Top*). By placing the diffuser in the optical path, the coherence of the beam is disrupted and the 2-D spatial pattern imposed by the DMD is accurately reconstructed in the sample only in a thin (axially-contained) image plane.

This image plane, optically conjugated with both the diffuser and the DMD chip comes into focus in the focal plane of the objective in the default 4f configuration such that the photo-stimulation and multiphoton imaging planes overlap axially (**Fig. 1c***, Middle*). Critically, translating the diffuser along the optical path causes its conjugate image plane in the sample to move as well, therefore axially decoupling the photo-stimulation plane from the focal (sample) plane of the imaging objective (**Figs. 1b-c**, *Bottom*, **S1a-b**).

### Optical specifications of *ADePT*

To characterize the spatial precision of *ADePT*, we registered the DMD field of stimulation (DMD pixel size = 2.4 µm at the sample) to the primary CCD camera image (CCD1 pixel size = 1.1 µm at sample; CCD 1300-QF, Vosskuhler) and the multiphoton field of imaging into a common reference frame (Methods). We verified that the targeting of DMD-modulated spatial light patterns was precise (< 5 µm error, fluorescence restricted to the microbeads targeted by the DMD photo-stimulation masks with minimal spillover to adjacent, off-target microbeads, **Figs. 1j**, **S1d**).

To determine the lateral and axial resolution of photo-stimulation setup, we projected light spots of increasing nominal diameters onto a glass coverslip spin-coated with 1µm thick rhodamine B layer (**Figs. 1e-g**). We measured the resulting fluorescence signals by imaging the rhodamine-coated coverslip from below with an inverted (secondary) microscope. The secondary microscope was set up so as to be confocal with the primary microscope objective (**Figs. 1d-g**, **S1a-d**, **S2**)^6^. Each spot was sampled through the fluorescent coverslip by moving the primary objective up or down in steps of 1-2 µm. We used the images acquired by the primary (CCD1) and secondary (CCD2) cameras (**Fig**. **S2a**; CCD1300QF, Vosskuhler) to obtain the lateral (x-y) and axial (z) extent of the light intensity profile at full width half maximum (FWHM, **Figs. 1d-i**). Using this strategy, we a projected grid pattern and measured the ensuing fluorescence. One pixel-wide lines could be easily resolved when 2 pixels (∼5 µm) part. Projecting circular spots, we measured a 6.2 ± 1.6 µm lateral FWHM (**Fig. 1d-e**). Below this size, decreasing the nominal spot size did not change the spot’s x-y FWHM, potentially due to light scattering from the extended spot, light loss and imperfections in the setup’s optical transfer function.

In the axial dimension, we observed a linear dependence between the axial and lateral extent of the light patterns projected (x-z FWHM vs. x-y FWHM of photo-stimulation spots, **Fig. 1i**), as reported previously for light patterning techniques such as digital holography^6,12,84^. As expected, the ratio between the axial vs. lateral resolution was inversely proportional to the diffuser divergence angle (slope = 1.52 when using a *grit ground* diffuser, #38-786 vs. 1.62 for a 10° *holographic* diffuser, #48-506 vs. 2.12 for a 5° *holographic* diffuser, #55-849, Edmund Optics, **Fig. 1i**). Note that this differs from the focusing regime of a Gaussian laser beam, where the axial extent of the focal spot increases quadratically with the lateral extent of the spot. To reduce excitation light loss, while maintaining reasonable axial resolution, we settled on a 5° *holographic* diffuser, a diffractive optical element designed to limit the spread of transmitted light to a specified solid angle (e.g., 5°, #55-849, Edmund Optics). We measured 51 ± 2 % light loss through the 5° *holographic* diffuser. Using the diffuser only slightly under-filled the angular aperture of the primary objective and did not affect the lateral resolution of the system. Across the field of stimulation, we measured a 34.5 ± 5.6 µm x-z FWHM.

Beyond determining axial resolution, the choice of diffuser also governs the axial decoupling of photo-stimulation and imaging planes. Translating the diffuser along the optical path causes its conjugate image plane in the sample to move as well, therefore axially decoupling the photo-stimulation plane from the focal (sample) plane of the imaging objective (**Figs. 1b-c**, Bottom, **S1a-b**). To calibrate the decoupling of photo-stimulation and imaging planes, we projected a checker-board light pattern on a rhodamine B coated coverslip. In the default configuration, the diffuser is conjugate to the sample (objective focal plane), the (secondary) objective of the inverted microscope is confocal with the primary objective, and the rhodamine coverslip is placed in their shared focal plane (also the sample plane). The photo-stimulated pattern thus appears in focus on both CCD1 (primary objective) and CCD2 (secondary objective). As we lowered the primary objective, as one would to image deeper structures in a sample, the projected pattern also moved lower (below the coverslip). This can be observed as a blurred image of the photo-stimulation pattern on CCD2 as its imaging plane is still conjugate to the rhodamine coverslip (**Fig. S2a**). We determined the extent of the diffuser displacement necessary to re-focus the projected pattern back onto the rhodamine coverslip by translating the diffuser away from the objective until a sharp image of the checker-board reappeared on CCD2 (**Fig**. **S2a**). Moving down the primary objective by incremental amounts (100 µm) and monitoring the required diffuser displacement via the secondary microscope, we obtained a calibration curve, describing the effective decoupling of the imaging and photo-stimulation planes up to 500 µm (i.e. spanning the 50 mm range of a Newport 433 translation stage we used to mount the diffuser, Methods, **Fig**. **S2b**). Over the axial displacement interval sampled, we observed a linear relationship between translating the objective axially and the compensatory displacement of the diffuser required to bring the pattern back into focus, proportional to the square of the effective optical magnification in our setup (theoretical vs. effective magnification ∼11.1 X vs. ∼9.3 X, **Fig. 1k**, Methods).

To characterize the propagation of DMD-generated light patterns through brain slices of different thickness^76,77^, we placed brain slices (Methods) atop a fluorescent coverslip, and recorded images of the projected patterns from underneath with the auxiliary microscope. As expected, the projected patterns were substantially impacted by light scattering, which constrains using this approach to photo-stimulating superficial structures within ∼100 µm from brain surface (**Fig. S3**).

### Mapping functional connectivity in the intact brain

*ADePT* enables investigating the logic of functional connectivity in layered neuronal circuits where input and output circuit elements do not necessarily lie in the same plane. In particular, our one-photon DMD based implementation is well-suited for understanding information flow in areas where sensory stimuli evoke structured spatiotemporal patterns of activity that extend over several 10s of micrometers (e.g. glomeruli in the olfactory bulb, whisker barrels in barrel cortex, etc.), and the input layer is close to the brain surface. As a proof of principle, we used *ADePT* to investigate the role of different local inhibitory and excitatory connections in shaping the output of the mouse olfactory bulb. Individual odors activate distinct spatial-temporal combinations of glomeruli. This odor-evoked activity is heavily reformatted within the OB by numerous excitatory and inhibitory interneurons and relayed to other brain areas in the form of activity patterns of the mitral/tufted cells. To date, understanding the OB computations has been challenging due to constraints in precisely and flexibly controlling the activity of individual input nodes (glomeruli) using traditional odor stimuli. Using *ADePT*, we can bypass these constraints by direct photo-stimulation while simultaneously monitoring the OB outputs at high throughput and cellular resolution.

### Application 1: Mapping glomerular receptive fields of sister mitral/tufted cells

We used *ADePT* to first identify cohorts of *sister* output cells in the OB – mitral and tufted cells that receive excitatory input from the same *parent* glomerulus^34,63–72^ – and then assess whether they integrate lateral inputs from different sets of glomeruli in the OB. Indeed, *sister* mitral and tufted cells extend multiple secondary (lateral) dendrites, which may each contact different sets of inhibitory interneurons^34,63,64,85^. Previous work has shown that while *sister* cells are generally responsive to the same set of odors, they are not redundant in their odor responses: pairs of *sister* cells are as different as pairs of *non-sisters* in their response time with respect to inhalation onset^34,63^. A potential mechanism for temporal differences in the odor-evoked responses of *sister* cells is differential inhibitory input from other glomeruli. However, to date, mapping glomerular receptive fields of individual mitral/tufted cells has been difficult given technical limitations in precise control of single-glomerulus activity in conjunction with flexible monitoring of responses at a high-enough yield to identify and compare cohorts of *sister* mitral/tufted cells. Using *ADePT*, we bypass the constraints imposed by odor stimulation and directly photo-stimulate individual glomeruli on the OB surface. In parallel, we monitor the responses of many mitral/tufted cells simultaneously at cellular resolution using multiphoton calcium imaging. To this end, we used mice expressing an opsin in the mature olfactory sensory neurons (OSN), OMP-Cre x ReaChR-mCitrine^86^ and the genetically encoded calcium indicator (GCaMP6f/s^87^) in the mitral and tufted cells (Thy1-GCaMP6s^88^; **Fig**. **S4a**, Methods).

We stimulated glomeruli by projecting DMD-modulated light masks on top of individual glomeruli, identified based on their baseline fluorescence (Methods). We used photo-stimulation spot sizes smaller (30-50 µm) than the targeted glomeruli (Avg. ∼75 µm)^34,89^. To assess the resultant responses in the mitral/tufted cells, we strobed between stimulating OSN axon terminals (50-150 ms stimulation) and monitoring calcium responses (5-10 Hz imaging rate) in either the postsynaptic mitral/tufted dendritic tufts in the glomerular layer (same z-plane) or the cell-bodies (100-270 µm below the photo-stimulation plane).

First, we calibrated the stimulation light intensity to ensure single glomerulus specificity – photo-stimulation of OSNs only within the targeted glomerulus while avoiding spurious excitation of passing-by OSN axons that terminate in neighboring glomeruli. To assess the efficacy of our approach, we set the photo-stimulation and multiphoton activity readout planes to overlap axially in the glomerular layer, such as to stimulate the OSNs and image the resultant postsynaptic glomerular response in the dendrites of MT cells. Photo-stimulation induced rapid and reliable changes in fluorescence (**Fig. 2**). For each photo-stimulation mask (**Figs. 2b-e**, **S4b**), we constructed a two-dimensional light activation map (2DLAM) at sub-glomerular resolution^34^ (e.g. 40 x 40 grid of 7.2 µm pixels, Methods). To ensure we operated in a regime of specific glomerular activation, we successively lowered the light intensity until no light-triggered activity was observed in neighboring glomeruli (15-to-2 mW / mm^2^, 122 glomeruli, 21 FOVs, 7 mice, **Figs. 2f**, **S4b-c**). Mitral and tufted dendritic GCaMP responses of individual glomeruli to direct photo-stimulation varied in amplitude, potentially as a function of strength of synaptic transmission of different glomeruli and intra-glomerular inhibition. Across mice, we identified a range of intensities that on average resulted in specific and reliable activation of targeted glomeruli and negligible activation of adjacent glomeruli (>90 % specificity, Methods, **Fig. 2f**). When only glomerular pixels are considered (**Fig. 2h**), ∼5-8 % of significant pixels were in off-target glomeruli, potentially due to spurious stimulation of fibers of passage.

**Figure 2.**
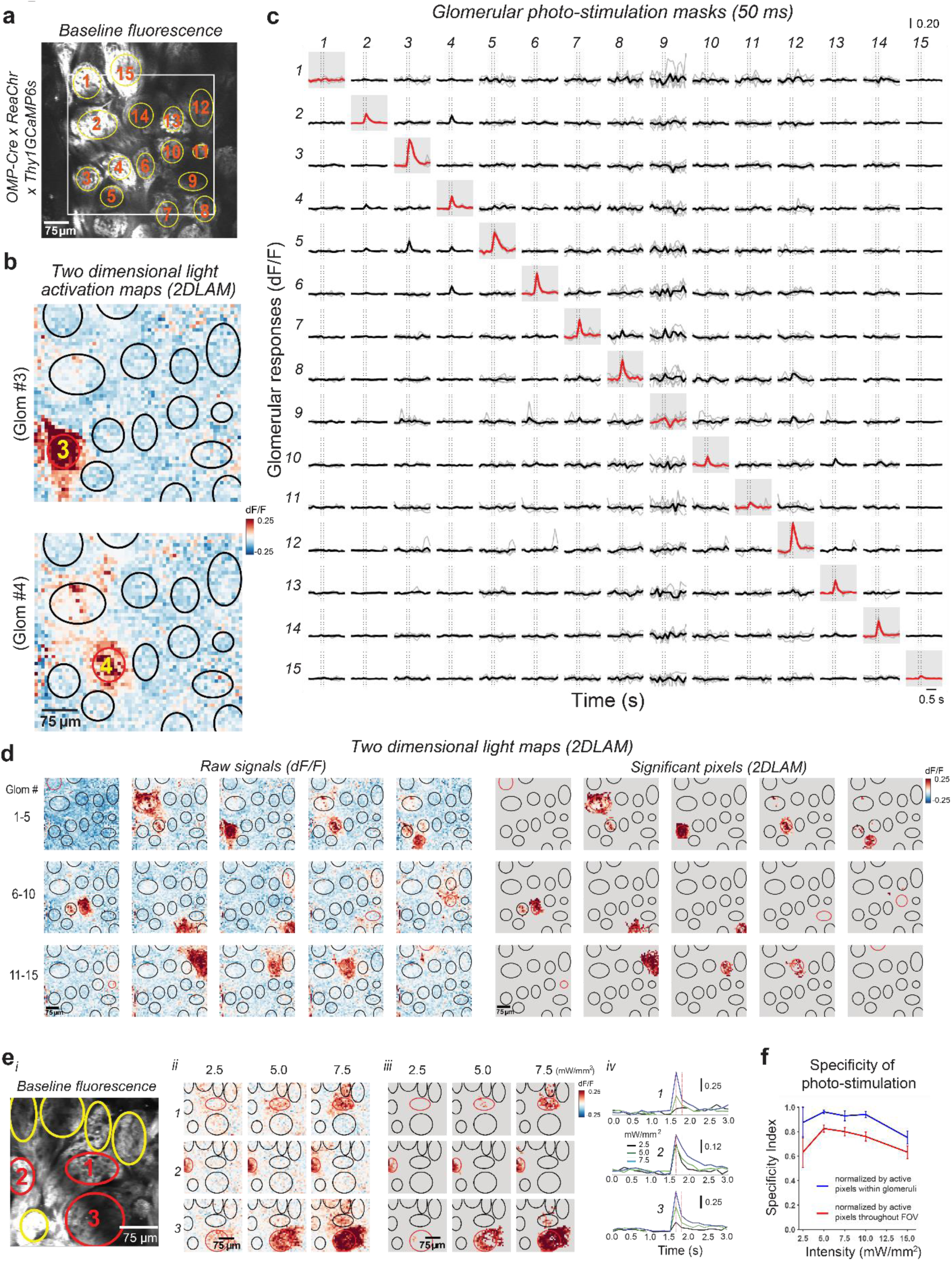
Targeting individual glomeruli by photo-stimulation of olfactory sensory neuron terminals and monitoring glomerular responses of sister mitral and tufted cell dendrites. **(a)** (i) Glomerular resting fluorescence in OMP-Cre x ReaChR-citrine x Thy1-GCaMP6s mice; **(b)** Fluorescence changes (two dimensional glomerular light activation maps, 2DLAMs) evoked by photo-stimulation of 2 example glomeruli (#3, #4) out of 15 glomeruli targeted in the example field of view (5.0 mW/mm^2^ for 50 ms). Red ellipses mark the location and contour of the projected light masks. Color scale in heat map indicates ΔF/F, comparing fluorescence after light stimulation to baseline; **(c)** Response matrix of fluorescence traces (individual repeats in gray, and mean signal as thicker lines) corresponding to 15 glomerular regions of interest versus the 15 (matching) glomerular light masks projected on the OB surface. Red traces mark responses of the targeted glomerulus in a given trial. **(d)** (Left) Fluorescence changes (two dimensional glomerular light activation maps, 2DLAMs; 5.0 mW/mm^2^) evoked by photo-stimulation of the 15 glomeruli targeted in the example field of view (**a**). (Right) Significant 2DLAMs. Non-significant responses were thresholded to gray background. Out of 15 targeted glomeruli, 5 glomeruli were considered non-responsive, given our signal threshold criteria (Methods). Of the 10 responsive light maps, 8 showed specific responses (pertaining to the target glomerulus), and 2 light maps were not specific (responses were observed across multiple glomeruli). **(e)** Example target glomeruli from a different example FOV across a range of three light intensities (2.5, 5.0 and 7.0 mW/mm^2^). Projected light masks match the red ellipses in (i) and (ii). Color scale in heat map indicates ΔF/F, comparing fluorescence triggered by light stimulation to baseline; (iii) Same as (ii) for statistically significant glomerular photo-stimulation responses (rectified 2DLAMs); (iv) Fluorescence traces (mean signal for each light intensity) corresponding to three regions of interest (ROI) in each heat map, whose position is indicated by red ellipses in i-iii. Different intensities are represented by different color traces (red - 2.5mW/mm^2^, green - 5.0mW/mm^2^, blue - 7.5mW/mm^2^). **(f)** Specificity of glomerular responses across light intensity. Blue trace: responsive pixels within the target glomerulus normalized by all responsive pixels located within glomerular boundaries across the FOV. Red trace: responsive pixels within the target glomerulus normalized by all responsive pixels within the FOV, including those spanning the juxtaglomerular space, surrounding glomeruli (Methods); N=7 mice, 21 FOVs; responsive vs. photo-stimulated glomeruli: 2.5mW/mm^2^, 12/74; 5.0mW/mm^2^, 48/122; 7.5mW/mm^2^, 27/57; 10.0mW/mm^2^, 48/98; 15.0mW/mm^2^, 48/77.

Next, we axially decoupled the photo-stimulation and imaging z-planes to monitor light-triggered responses in the cell bodies of the corresponding *daughter* M/T cells (receiving excitatory input from the targeted glomeruli, and 100-270 µm deep from surface), while still photo-stimulating individual glomeruli on the bulb surface (**Figs. 3a-b**, **S5a-m**). Photo-stimulation of individual glomeruli evoked robust excitatory responses in ∼19 % of the cells (162/856). The majority of these responsive cells (141/162) were activated by only one unique glomerulus that we ascribed as their corresponding parent glomerulus. A small proportion of cells were activated by more than one glomerulus and excluded from were further analysis (21/162 cells, **Fig. 3b** *iii*, cell 30). By comparing the identity of parent glomeruli across the imaged M/T cells, we identified cohorts of *sisters cells* that were activated by different parent glomeruli. On average, across experiments (10 FOVs, 6 mice, 27/46 response evoking glomeruli) we identified 6.9 ± 6.0 (Avg. ± SD) *sister* mitral and tufted cells per glomerulus (4 SD response threshold, see Methods, **Figs. 3c**, **S5a**).

**Figure 3.**
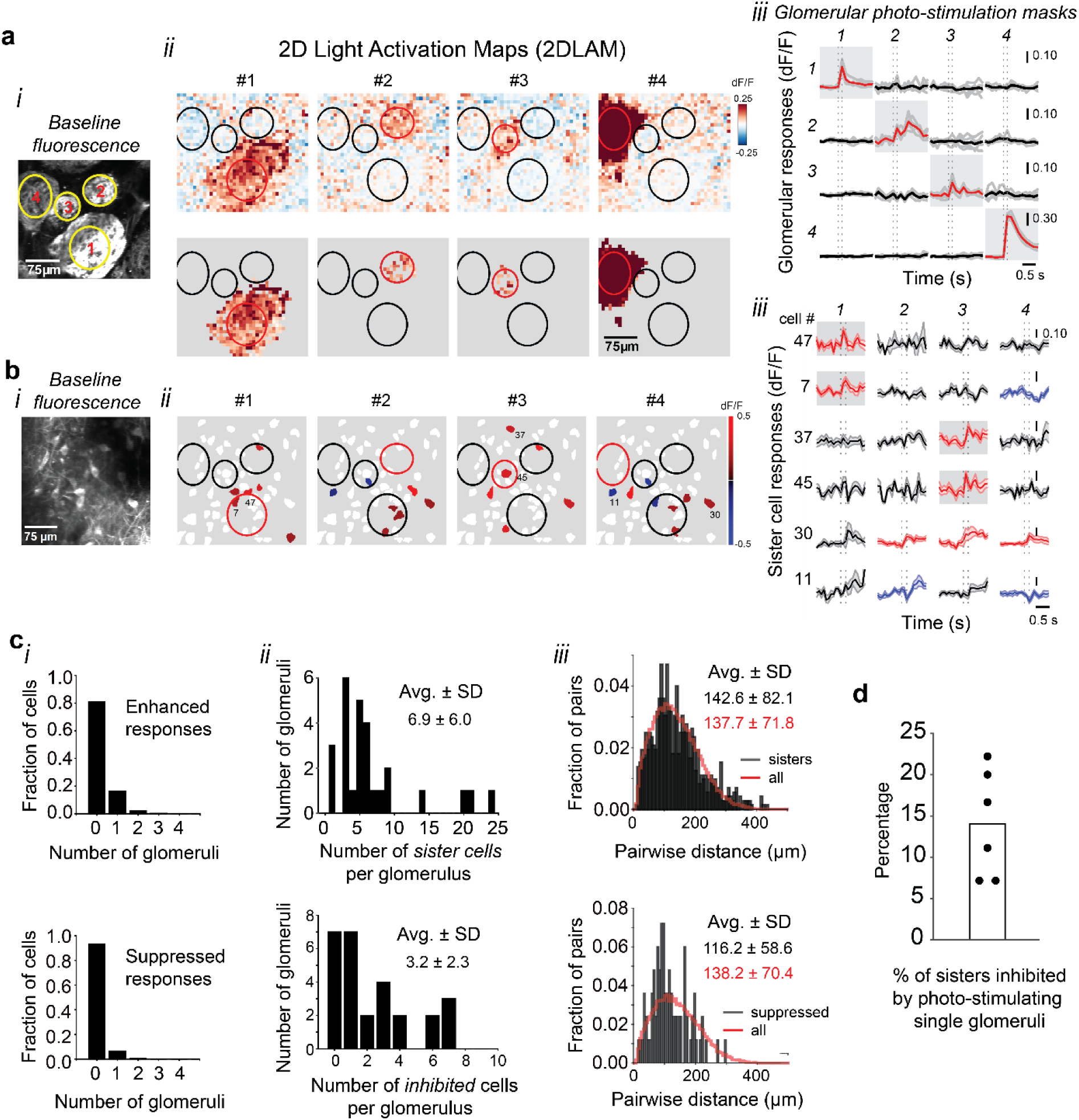
Identifying sister mitral and tufted cells via axially decoupled glomerular stimulation and two photon fluorescence imaging. anaestherized (**a-b**); awake+anaesthetized (**c-d**);**(a)** (i) Resting glomerular fluorescence (ReaChR-mCitrine and GCaMP6s); (ii) Fluorescence changes (raw, Top and significant pixels glomerular 2DLAMs, Bottom) evoked by photo-stimulation of four target glomeruli (7.5 mW/mm^2^); (iii) Response matrix of fluorescence change traces (individual repeats in gray, and mean signal as thicker lines) corresponding to four glomerular regions of interest versus the four corresponding (matching) glomerular light masks projected on the OB surface. Red shaded areas mark responses of the targeted glomerulus in a given trial. (**b**) (i) Resting fluorescence (GCaMP6s) of mitral cell bodies at the same x-y coordinates, 210 µm below the glomerular field of view in (**a**); (ii) Significant cell body responses evoked by photo-stimulation of the four target glomeruli (**a**, 7.5 mW/mm^2^); (iii) Average fluorescence response traces corresponding to six example mitral cells in each heat map, whose positions are indicated by numbers in (ii); shaded area marks SEM. Two example pairs of sister cells (47, 7 with respect to parent glomerulus #1; and 37, 45 with respect to parent glomerulus #3) are shown. Note that cell #30 is non-specifically responding to photo-stimulation of multiple glomeruli (#2,#3,#4). In addition, glomerular photo-stimulation triggered sparse suppressed responses (e.g. cells 7,11) intermingled with the sister cells. **(c)** (i) (Top) Specificity of enhanced responding mitral and tufted cells as a function of the targeted glomerulus (N = 6 mice; 27 responsive and specific glomeruli / 46 photo-stimulated target glomeruli; 856 mitral & tufted cells). Note that most cells were non-responsive, or responded specifically to the photo-stimulation of the target glomerulus (13 % of cells had off-target responses to more than one glomerulus; also see **Fig. S5a**, Methods); (Bottom) Same for suppressed responses. (ii) (Top) Number of observed sister cells (enhanced responses) per glomerulus photo-stimulation across fields of view (Left; 6.9 ± 6.0, N = 27 FOVs); (Bottom) Number of cells per glomerulus whose baseline fluorescence was suppressed upon photo-stimulation of the same glomeruli (Right; 3.2 ± 2.3, N = 20 FOVs); (iii) (Top) Pairwise distance between sister cells vs. between any cells in the field of view (142.6 ± 82.1 µm vs. 137.7 ± 71.8 µm; 725 vs. 29,850 pairs). (Bottom) Pairwise distance between suppressed cells vs. any cells in the field of view (116.2 ± 58.6 µm vs. 138.2 ± 70.4 µm, 83 vs. 19,607 pairs). Unless specified Avg. ± SD values are shown. **(d)** Photo-stimulation of individual glomeruli triggered inhibitory responses in only a small subset of the sister cells of a given identified cohort (Avg. ± SEM, 14.0 ± 2.7 %, N=6).

Consistent with previous literature^34,64,66,71,90^, *sister* cells were spread across the field of view and spatially intermingled with cells receiving inputs from other glomeruli (Avg. ± SD: 142.6 ± 82.1 µm pairwise distance between *sister* cells vs. 137.7 ± 71.8 µm between any cells in the FOV, **Figs. 3c**, **S5a**). Note that the statistics reported here underestimate the number of *sister* cells per glomerulus^66^ simply because of the limited number of different depths (z-optical planes) and the size of fields of view sampled for this experiment.

In addition to light-triggered excitatory responses, we also observed sparse light-induced suppression of the baseline fluorescence in 3.2 ± 2.3 cells per glomerulus (N = 20 FOVs). These suppressed cells were intermingled with the identified *sister* cells, and peppered the fields of view (64 suppressed cells; 3 SD response-to-baseline signal threshold; 116.2 ± 58.6 µm pairwise distance of suppressed cells vs. 138.2 ± 70.4 µm between any cells in the field). We interpret the observed decrease in M/T resting fluorescence as the suppression of respiration-locked baseline firing patterns^91–93^ due to lateral inhibition from the photo-stimulated glomeruli acting via local bulbar circuits. Interestingly, we could also observe such light-triggered suppression for a subset of the *sister* cell cohorts (6/27 *sister* cell cohorts, each associated with a different glomerulus) identified via *ADePT*. Photo-stimulation of individual glomeruli inhibited a small fraction of *sisters* within a cohort (1-2 *sisters* per cohort; Avg. ± SEM, 14.0 ± 2.7 %, N=6, **Fig. 3c**, **S5a**, Methods). These data suggest that *sister* cells, while sharing excitatory drive from the same parent glomerulus, receive sparse inhibitory inputs from different sets of glomeruli. This is in agreement with previous reports of differences in *sister* cells spike-timing across the respiration cycle^34,63,85^. As suppressed responses are small in amplitude, quantification of the sparseness and diversity of the inhibitory inputs of sister cells requires further systematic analysis by pairing odor stimulation to boost sisters’ responses with photo-stimulation of specific combinations of other (*non-parent*) glomeruli. While additional investigation is needed to understand the logic of these lateral inhibitory interactions, these experiments show that *ADePT* is well-suited for direct mapping of excitatory/inhibitory connections in layered neuronal circuits.

### Application 2: Assessing the spatial heterogeneity of long-range olfactory bulb inhibition

Next, we used *ADePT* to assess the spatial heterogeneity in the impact of GABAergic/dopaminergic (DAT+ cells, historically known as superficial short axon cells), a class of long-range inhibitory interneurons in the glomerular layer, on the activity of mitral and tufted cells. DAT+ neurons integrate excitatory inputs within individual glomeruli and, contrary to their ‘short axon’ namesake, establish chemical and electrical synapses with other interneurons and mitral/tufted cells within other glomeruli located as far as 1.5 mm away^41,75,76,80–82,94^. Given these long-range processes, DAT+ cells are thought to re-distribute focal glomerular odor inputs and convey long-range signals across the glomerular layer. Work from several groups including ours^40,41,75–82,95^ suggested that DAT+ neurons enable olfactory circuits to negotiate wide variations in odor concentration by implementing a form of gain-control (divisive normalization)^40,41,77,96^. However, it remains unclear whether DAT+ cells broadcast inhibitory inputs uniformly within a local neighborhood, or act selectively on OB output cells that share specific features such as similarity in odor tuning^41,76,95,97^. This is due to previous technical constraints in controlling the activity of DAT+ cells in conjunction with simultaneous imaging of activity in the mitral and tufted cells. We used *ADePT* to photo-stimulate DAT+ cells associated with individual glomeruli while monitoring their impact on either the spontaneous or odor-evoked responses of mitral/tufted cells.

We first tested the specificity of targeting DAT+ interneurons across glomerular regions of interest using mice co-expressing both an opsin (ChR2-mCherry^98^) and a calcium indicator (GCaMP6f^87^) in the DAT+ cells (**Fig**. **S6a**). We selected individual adjacent glomeruli for photo-activation, triggering and reading out DAT+ cell activity in the same optical z-plane. Through these experiments, we identified a range of light intensities across which the stimulus-responsive area remained confined to the glomerulus targeted for photo-stimulation (avg. spatial width constants of 40-60 µm; **Figs. 4a-c**, **S6b**, Methods). To assess the impact of glomerular DAT+ interneurons on the activity of mitral and tufted cells, we used mice expressing an excitatory opsin in DAT+ cells and the calcium indicator GCAMP6f in mitral/tufted cells (DAT-Cre x Thy1GCaMP6f^87,99^ mice injected retro-orbitally with ChrimsonR-tdtomato AAV-PHP.eB:Syn-FLEX-rc[ChrimsonR-tdtomato]^100,101^, **Fig**. **S7a**, Methods).

**Figure 4.**
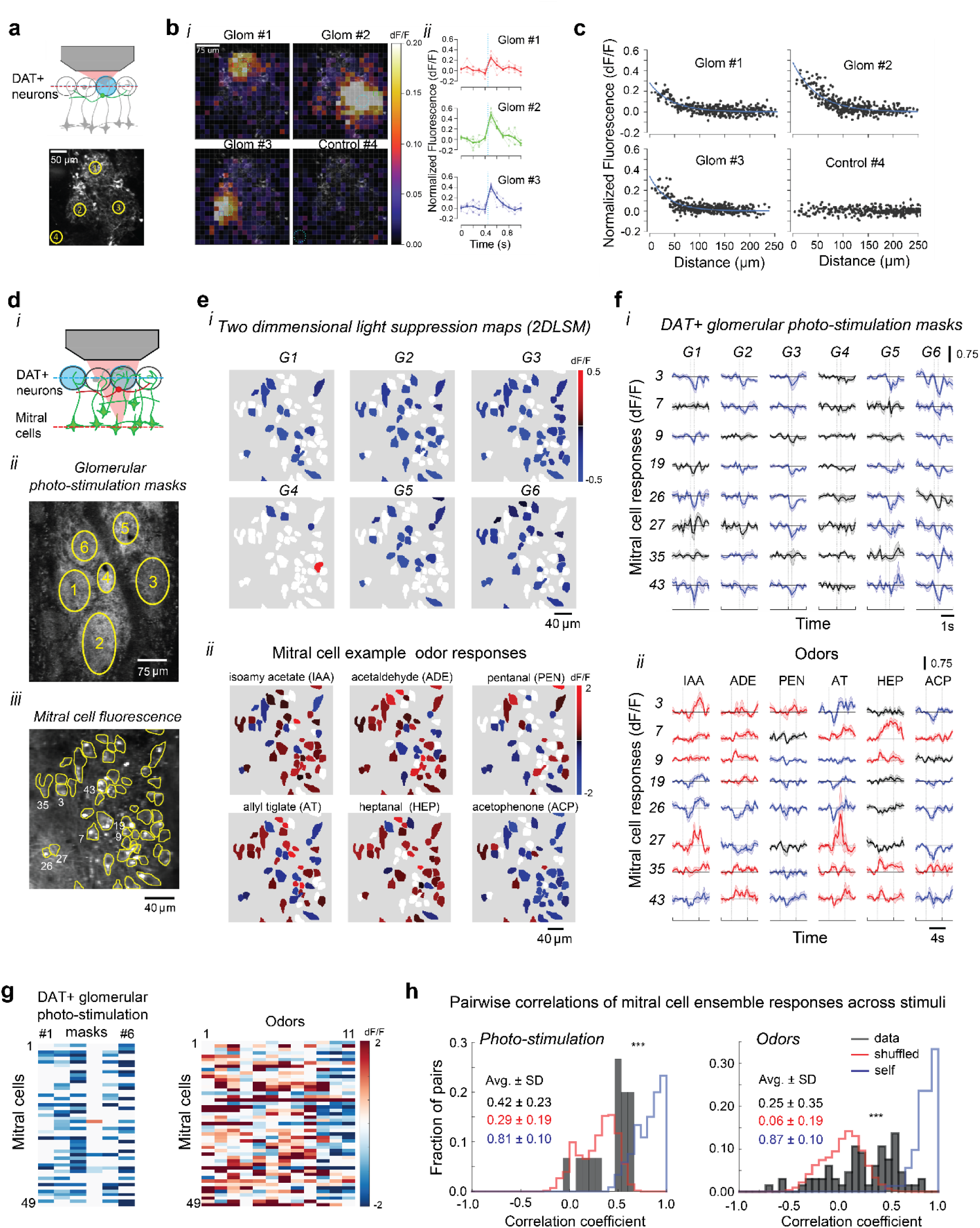
Glomerular photo-stimulation of DAT+ interneurons triggers dense, but spatial heterogeneous inhibition of mitral cell activity. **(a)** Schematic of in vivo experiment (Top) and baseline 2p fluorescence of three adjacent glomeruli in the field of view (Bottom); GABAergic/dopaminergic superficial short axon cells (DAT+) co-expressing GCaMP6f and ChR2-mCherry; circles and numbers indicate the location of projected light masks. **(b)** (i) Fluorescence (two-dimensional light activity maps) evoked by photo-stimulating DAT+ cells in glomeruli (1-3) and one (no apparent opsin expression) control site (4) marked in (**a**) using light masks equivalent to the dotted blue circles. Color scale in heatmap indicates ΔF/F, comparing fluorescence after light stimulation to baseline. (ii) Fluorescence traces (average, thick line, and four individual repeats, thin lines) corresponding to a region of interest (ROI) in each heatmap, whose position is indicated by matching color squares in (i). As in the case of OSN terminals photo-stimulation experiments, we flashed single 100 ms sub-glomerular light masks (20 µm diameter) between the imaging frames and recorded calcium transients in the DAT+ cells via multiphoton imaging; for analysis, the field of view was divided into a 20 by 20 grid (15 x 15 µm grid units), and the response was defined as the average change in fluorescence in each unit of the grid during the two frames (10 Hz) immediately following the light stimulus. **(c)** For each ROI (1-4), the evoked ΔF/F response is plotted against the distance of that ROI to the center of each stimulation pattern across repeats. For all stimuli except the control (4), the relationship was modeled as an exponential decay. **(d)** (i) Cartoon schematics: photo-stimulation of opsin expressing DAT+ cells while monitoring mitral cell activity (GCaMP6f) by axial decoupling of the photo-stimulation and imaging planes (awake). (ii) Example glomerular field of stimulation; mice express GCaMP6f in mitral and tufted cell glomerular dendritic tufts and ChrimsonR-tdtomato in DAT+ cells (not shown). (iii) Example field of view of mitral cells imaged while photo-stimulating DAT+ cells (200 ms, 7.5 mW/mm^2^) in the glomerular layer. **(e)** (i) (Top) Fluorescence changes (two dimensional light suppression maps) in mitral cells evoked by photo-stimulation of DAT+ cells across six glomeruli (G1-G6); (Bottom) Diversity of same mitral cell responses to six example odors; each cell in the field of view is displayed on a blue-to-red color scale indicating its response amplitude to light/odor stimulation; non-significant responses were thresholded to white (Methods). **(f)** (i) (Top) Dense, but heterogeneous inhibition of baseline fluorescence in example mitral cells across the field of view evoked by photo-stimulation of DAT+ cells across six different glomerular light masks (G1-G6); (Bottom) Odor responses of the same example mitral cells to six odors (isoamyl acetate, IAA; acetaldehyde, ADE; valeraldehyde, PEN; allyl tiglate, AT; heptanal, HEP; acetophenone, ACP). **(g)** Peak fluorescence changes (ΔF/F) of same mitral cells (49) across the field of view in response to glomerular DAT+ cells photo-stimulation (Left) and to a panel of eleven odors (Right). **(h)** (Left) Histogram of correlation coefficients of mitral cell ensemble responses across different pairs of DAT+ cell glomerular photo-stimulation masks (G1-G6). (Right) Same, across pairs of odors in the panel (1-11). Histograms of shuffled mitral cell index correlation distributions are shown in red; ‘self’ controls in blue (correlations between mitral ensemble responses to repeats of the same DAT+ photo-stimulation mask, Methods).

We photo-activated DAT+ neurons associated with individual glomeruli and monitored the tufted and mitral cell responses 120-275 µm below the photo-stimulated glomeruli (**Figs**. **4**,**5**). On average, glomerular DAT+ photo-stimulation targeted to *single* glomeruli resulted in dense, but *heterogeneous suppression* of the baseline fluorescence in mitral cells across the field of view (fraction of mitral cells modulated in example field of view: 0.40 ± 0.19; 200 x 200 µm, 294 cell-mask pairs, **Figs. 4d-h**, **S7b**). Stimulation of DAT+ cells associated with *different* glomeruli resulted in different patterns of mitral cells suppression (Avg. DAT+ masks pairwise correlation in ensuing MC responses = 0.42 ± 0.23; **Figs. 4e-h**, *Left*). The mitral ensemble responses to direct DAT+ photo-stimulation of *different* glomerular masks were more correlated than the average pairwise responses for controls where cell identities were shuffled (**Fig. 4h**, *Left*; 0.42 ± 0.23 vs. 0.29 ± 0.19, p<0.001, Wilcoxon signed-rank test) or across pairs of odors (**Fig. 4h**, *Right*; 0.42 ± 0.23 vs. 0.25 ± 0.35, p<0.001, Wilcoxon signed-rank test). As reference, we also calculated a ‘self’ correlation to assess the variability of photo-stimulation responses to the *same* DAT+ glomerular mask across trials (Methods). Responses of mitral cell ensembles across different DAT+ photo-stimulation masks were substantially less correlated than the ‘self’ control (0.42 ± 0.23 vs. 0.81 ± 0.14, p<0.001, Wilcoxon signed-rank test, **Fig. 4h**). These data demonstrate the significant impact of DAT+ neurons associated with different glomeruli on decorrelating the OB outputs.

**Figure 5.**
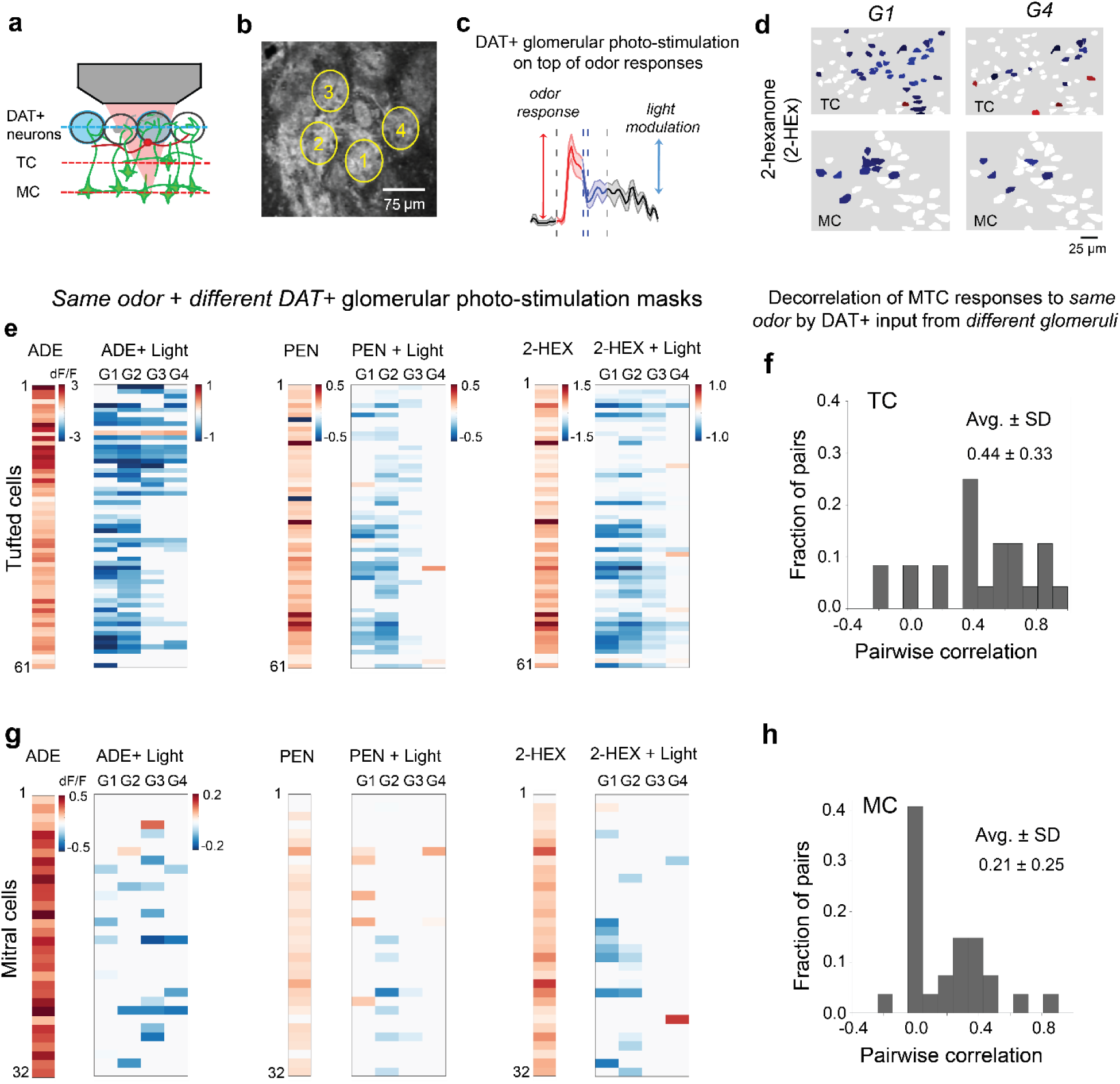
Photo-stimulation of DAT+ interneurons from different glomeruli on top of odor responses triggers decorrelates the mitral and tufted cell responses. **(a)** Schematic of the experiment: in vivo photo-stimulation of ChrimsonR-tdtomato expressing DAT+ cells, while monitoring its impact on mitral and tufted cell odor responses (GCaMP6f). **(b)** Baseline 2p GCAMP6f fluorescence of an example field of DAT+ stimulation in the glomerular layer. **(c)** Fluorescence change (dF/F) elicited by odor presentation in one example tufted cell, and its subsequent modulation by projecting of a glomerular light mask to boost DAT+ cell activity (500 ms, 10mW/mm^2^; 2s from odor onset). **(d)** Odor response modulation (dF/F) via boosting DAT+ cells associated with different glomeruli. Two dimensional light suppression/activation maps) of tufted (Top) and mitral cell (Bottom) responses to one example odor (2-hexanone, 2-HEX) evoked by photo-stimulation of DAT+ cells across 2 example glomeruli (G1,G4) from **(b)**. **(e**, **g)** Mitral (**e**) and tufted (**g**) cell peak odor responses (ADE, PEN, 2-HEX) and their modulation by DAT+ cell light masks associated with four different glomeruli (G1-G4) from the example field of view in **(b)**; also see **Fig**. **S8**. **(f**, **h)** Decorrelation of MTC responses to same odor by boosting DAT+ inputs from different glomeruli. Histograms of pairwise correlations of modulating the tufted cell ensemble responses (**f**) and of mitral cell ensemble responses (**h**).

Stimulation of DAT+ neurons results predominantly in the suppression of mitral and tufted cells from their baseline activity, whose interpretation may suffer from floor effects. To boost the baseline M/T cell fluorescence and further assess the impact of DAT+ neurons, we presented odors (4s) in conjunction with DAT+ cell photo-activation in the middle of the odor presentation period. Similar to the optogenetic stimulation only experiments described above (**Figs. 4d-h**), photo-stimulating DAT+ cells across *different glomeruli* during odor presentation also resulted overall in different patterns of suppression of the OB outputs in a given field of view (**Figs. 5a-h**, **S8b-d** TC: 0.44 ± 0.33; MC: 0.21±0.25; pairwise correlation of cell ensembles, p<0.001, Wilcoxon signed-rank test; modulation by light on top of odor: 78.0 % unmodulated, 18.2% suppressed vs. 3.8% enhanced MC-odor-pairs; N=2,214 pairs; and 54% unmodulated, 42.6% suppressed vs. 3.4% enhanced TC-odor pairs, N = 945 pairs). Consistent with previous reports, the activation of DAT+ neurons suppressed and decorrelated the mitral cell ensemble responses to a larger degree compared to the tufted cells.

Altogether, we find that the impact of DAT+ cells on the OB output is dense, but locally heterogeneous (non-uniform) with different mitral and tufted cells exhibiting diverse sensitivities to DAT+ inhibition originating from a given glomerulus (**Figs**. **4**,**5**). Further investigation will determine whether DAT+ cell action shapes the MC and TC cell odor responses in a systematic manner as a function of similarity in their odor responses.

In summary, we show that *ADePT* can be flexibly combined with genetic strategies and/or sensory stimuli to probe the contribution of both excitatory and inhibitory neuronal types in shaping the output of optically accessible neuronal circuits.

## Discussion

We demonstrate a readily implementable method (*ADePT*) for optical control and recording of activity in three-dimensional neural structures. *ADePT* allows arbitrary axial shifting of 2D photo-stimulation light patterns of choice relative to the focal plane of the imaging objective by translating a motorized diffuser along the photo-stimulation optical path (**Fig. 1**), and builds on work from other groups to constrain DMD patterns using diffusers^38,48,49^. Using *ADePT*, we were able to maintain precise spatial control over a range of activation powers and demonstrated the usefulness of the system by mapping both excitatory and inhibitory functional connectivity in the mammalian olfactory bulb.

*ADePT* occupies a distinct niche among existing strategies for axially-decoupled photo-stimulation and imaging. It differs from other strategies of pairing DMD-based photo-stimulation and multiphoton imaging in its flexibility of adjusting the depth of imaging with respect to the photo-stimulation^46^. Several approaches have been developed over the past two decades to target multiple optical z-planes simultaneously, or in quick succession, each with characteristic trade-offs in resolution, field of view, cost, and complexity. Electrically tunable lenses (ETLs) offer a simple and low-cost route to axial decoupling, but may introduce spherical aberrations that deteriorate the point spread function and limit the usable field of stimulation^20–22^. AOD-based arbitrary access systems afford rapid and precise 3-D targeting, but are technically demanding, cost-prohibitive, and the size of the field of photo-stimulation is constrained by steep power requirements^15,24–28^. Digital holography combined with temporal focusing achieves the highest spatial precision – including single-cell resolution – but shares similar cost and complexity limitations, and likewise scales poorly to large fields of stimulation^8,9,12,28–33^. Compared to these approaches, *ADePT* enables optogenetic control across a wide field of view without aberrated imaging, and is straightforward to implement as a modular add-on to an existing two-photon microscope at relatively low cost. The principal trade-off is resolution: in its current implementation, *ADePT* does not afford the precision to target individual cell bodies, but is highly effective for controlling activity at the level of larger functional units – such as olfactory glomeruli or whisker barrels – while simultaneously monitoring downstream cellular responses at high throughput via multiphoton imaging. A second constraint is penetration depth: as a one-photon approach, *ADePT* is subject to light scattering and is best suited to superficial brain layers, within 100 µm from surface. Together, these trade-offs position *ADePT* as a practical tool for circuit-level functional mapping in optically accessible brain regions, particularly where large-scale patterned stimulation and cellular-resolution readout are required in tandem.

In proof-of-principle experiments, we used *ADePT* to investigate the functional connectivity of olfactory bulb circuits in the mouse. We controlled optogenetically the activity of select excitatory (**Figs. 2**,**3**) and inhibitory (**Figs. 4**,**5**) cell types in the input layer (OSNs vs. GABAergic / dopaminergic DAT+ interneurons) at *sub-glomerular* resolution, while monitoring their impact on the responses of the principal neurons (mitral and tufted cells). Our approach allows discriminating between different inhibitory inter-glomerular connectivity schemas (e.g. uniform vs. heterogeneous). Specifically, we found that DAT+ cells associated with different glomeruli mediate dense, but heterogeneous (non-uniform) suppression of MC and TC baseline and odor responses (**Figs**. **4**,**5**). By targeting optically addressable glomeruli (**Figs**. **2**,**3**), we identified multiple cohorts of daughter *sister* mitral and tufted cells driven by different *parent* glomeruli. For each cohort of sister cells analyzed, only a small subset was suppressed by photo-stimulation of individual (*non-parent*) glomeruli, suggesting as previously proposed^34,63,64^, that sister cells receive inhibitory input from different sets of glomeruli. Because responses were often weak and baseline noisy potentially due to spontaneous activity, caution needs to be taken in interpreting these results, as some of suppressed responses may go undetected. Pairing of glomerular photo-stimulation with odor presentation to boost sister mitral and tufted cell responses will enable assessing the sparseness and diversity of inhibitory inputs of sisters. In addition, systematic stimulating specific spatial-temporal combinations of multiple glomeruli in conjunction with monitoring of fast response dynamics, will provide insight on the linearity of summation effects, and how inhibitory connectivity relates to the degree of similarity in the odor response tuning of the interacting glomeruli.

More broadly, simultaneous patterned photo-stimulation and multiphoton imaging in different axial planes enables mapping functional connectivity not only in the olfactory bulb, but can be implemented between any optically accessible brain regions (across layers) including the neocortex. For example, *ADePT* is well positioned for mapping functional connectivity in the barrel cortex within and across barrels^102,103^, as well as bidirectional interactions between cortical visual areas (V1 vs. PM, AL, etc.)^104,105^, visuomotor (M2-V1)^106–108^ or audiomotor circuits (M2-A1)^109,110^, etc. Given its relative simplicity, cost-effectiveness, and ease-of-use, compared to other axially decoupling optical imaging and photo-stimulation strategies, *ADePT* is readily applicable for investigating the logic of neural connectivity.

## Acknowledgements

The authors acknowledge R. Eifert and D. Cowan for technical support. This work was supported by NIH 5R01DC012853-05, NSF IOS-1656830 to D.F.A. and Beckman Institute CLOVER (CLARITY, Optogenetics, and Vector Engineering Research) to V.G.

## Data and code availability

All data matrices included in the analyses presented here representing mitral/tufted and GABAergic/dopaminergic DAT+ interneuron GCaMP responses triggered by odor and optogenetic light stimuli, as well as the code used for the instrument control and analysis are available upon request.

## Authors contributions

conceptualization, investigation, methodology, formal analysis, visualization, software, data curation and writing, M.K., F.A.; methodology, formal analyses, visualization, data curation, validation, software, S.K.K., S.S.T; methodology, visualization, validation, D.E.H., W.G.B., P.S.V., H.C, M.B.D.; reagents development, Z.Q., V.G.; conceptualization, methodology, formal analysis, visualization, and writing, A.B.; conceptualization, methodology, formal analysis, visualization, software and writing, P.G.; conceptualization, methodology, investigation, methodology, writing, supervision, project administration, and funding acquisition, D.F.A.

## Methods

### Combined two-photon imaging and photo-stimulation microscope

For *two photon imaging* (**Fig. 1b**), the output from a Ti:Sapphire laser (Coherent Chameleon Ultra II) was magnified by a 1.5x beam expander (L1, L2, focal lengths, FL: 40, 60 mm), then reflected off two mirrors (10D20ER.4, Newport) and a periscope onto a galvanometric mirror scanning system (GM, 6215HB, Cambridge Technologies). The semi-stable stationary point between the GM was expanded and imaged to fill the back aperture of the microscope objective by a telescope (L3, AC300-050-B, L4, AC508-300-B, FL 50, 300 mm, Thorlabs). The objective and microscope stage were mounted on motorized linear translators (x-y stage: 462-XY-M; z-stage: CMA-25CCCL, Newport) connected to a three-axis motion controller (Newport ESP301). Fluorescence light from the sample was collected by the objective, then reflected by a dichroic mirror (DM1, 680dcxr, 50 mm diameter, Chroma) placed atop of the imaging objective (Olympus 20x 1.00 NA XLUMPlanFLN) and a second short-pass dichroic mirror (DM2, 505 nm, FF505-SDi01-50.8-D, 50 mm diameter, Semrock) towards a GaAsP PMT (H11526-20-NF, Hamamatsu) for green or red fluorescent detection (sliding holder, LCFH2 green, FF03-525/50-25 and red, FF01-607/70-25 emission filters, Semrock). The back aperture of the microscope objective was imaged into the PMT aperture by a telescope (L5/L6, FL: 100/32 mm; L5 – LA4545-A (UV fused silica); L6 - AL4532-A, Thorlabs). The current output of the PMT was transformed to voltage, amplified (SR570, Stanford Instruments) and digitized using a data acquisition board (NI-PCI-6115, National Instruments). The same board was also used to generate signals for controlling the galvano-metric mirrors. By rotating DM2 by 90° on a custom holder (CSHL, Machine Shop), light was directed either to the PMT or to a CCD camera (Vosskuhler 1300-QF, **Fig. 1b**). The image of the sample was further minified by a telescope (L7/L8, focal lengths: 100/40 mm) and captured on the CCD chip (CCD 1), allowing for a larger field of view of approximately 1,500 x 1,200 µm. Additional short pass filters (FF01-750/SP-50, 50 mm diameter, Semrock) were used in front of light detectors (CCD and PMTs) to block stray light from the infrared multiphoton excitation laser.

For *patterned photo-stimulation*, the beam from a 488-nm laser (Coherent Sapphire 488, 200mW) was magnified using a beam expander (L12/L11, FL: 30/300 mm) and reflected by a mirror and a periscope (RS99, Thorlabs) onto the surface of a DMD chip (DMD 0.7” XGA VIS, Vialux GmbH, 1024 x 768 pixels, 13.68 µm micro-mirror pitch). The DMD chip was controlled using a Vialux DLP-Discovery 4100 board and the corresponding application programming interface (Vialux GmbH). To obtain direct access to the chip, we removed the projecting lens and prism installed in front of the DMD by the manufacturer. The steep angle of incidence of the photo-stimulation beam onto the DMD was carefully tuned to ensure maximal diffraction efficiency. The patterned beam reflected off the DMD was further magnified by a L10/L9 telescope (FL 100/200 mm), and coupled into the microscope by DM1. An iris (SM2D25, Thorlabs) was placed at the front focal plane of L10 to block the higher diffraction order replicas of the DMD-generated spatial pattern. The photo-stimulation pattern was further projected into the sample by the tube lens (L5, FL 100 mm) and the principal objective (Olympus 20x 1.00 NA XLUMPlanFLN).

### Diffuser

A diffuser was used to axially confine the photo-stimulation pattern. The diffuser was mounted on a translational stage (Newport, 433-series, #BZA647202) connected to a stepper motor. The motor was controlled using an open source Arduino-based system^111^. In the home position (imaging and photo-stimulation planes superimposed), the diffuser sits at the focal plane of the tube lens L5. Three different round diffusers (25 mm diameter) were tested, as detailed in the text: two holographic diffusers with 5° and 10° divergence angle (5° - #55-849, 10° - #48-506, Edmund Optics), and a grit ground glass diffuser (#38-786, Edmund Optics). Two fast shutters (Uniblitz VS14) were placed in front of the PMT and the blue photo-stimulation laser respectively, and operated in anti-phase so that only one shutter was open at a time such as to alternate between periods of photo-stimulation and photon collection.

### Excitation power control and lenses

The excitation two photon laser power was controlled by a Pockels cell (Conoptics 350-80 BK) placed in the imaging path. An additional filter (775 nm long pass, 25 mm diameter, #86-069, Edmund Optics) was added to the excitation path to block a residual red component from the Ti:Sapphire laser, that would otherwise saturate the red PMT. When required, additional neutral density filters (Thorlabs, NDK01) ranging from ND 0.2 to ND 2.0 were added to the photo-stimulation path to further reduce the power. All lenses (Thorlabs) had anti-reflective coating for either infrared (imaging path) or visible (photo-stimulation and detection paths) light. A UV fused silica lens was used instead of the more common N-BK7 lenses in position L5 (LA4545-A) to avoid spurious phosphorescence signals following illumination with intense blue light pulses (see *Phosphorescence Artifacts*).

### Auxiliary microscope

An auxiliary microscope was used during characterization experiments, as described in *Lutz et al. (2008)*^6^. We placed a secondary objective, tube lens and green emission filter (HQ510DCLP, Chroma) below the main imaging objective to image the focal plane of the main objective onto a CCD camera (Vosskuhler 1300-QF, **Fig**. **S2**). Depending on the experiment, different objectives (Olympus 4X, 0.13 NA UPlanFlN or Olympus 20X, 0.50 NA UPlanFlN) were used in combination with a 150 mm focal length tube lens to obtain either a minified image of the entire field of photo-stimulation or a higher resolution image of a smaller field. The microscope framework was constructed with a Thorlabs 2” cage system. The secondary objective was held either by a kinematic mount for fine tilt adjustment (KC2-T, Thorlabs), or by a laterally-translatable platform. The photo-stimulation patterns were imaged onto a coverslip spin-coated with a thin (∼1 µm) rhodamine B layer (courtesy of V. Emiliani, Paris Descartes University).

### Calibrating decoupling of photo-stimulation and imaging planes

We used the auxiliary microscope to calibrate the decoupling of photo-stimulation and imaging planes (**Fig**. **S2a**). Initially, the diffuser was positioned at a plane conjugated to the focal plane of the principal and auxiliary objectives, and the photo-stimulation pattern was in focus on the coverslip, appearing sharp both on the principal (primary) and the auxiliary (secondary) CCD cameras (**Fig**. **S2a**, *Left*). The principal objective was then moved closer to the coverslip to simulate shifting the imaging plane to deeper layers in a biological preparation. At this point, the photo-stimulation pattern was still imaged (projected into sharp focus) on the focal plane of the principal microscope objective (now below the coverslip), and no longer in focus on the secondary CCD camera. To move the photo-stimulation pattern back to the initial position, we moved the diffuser away from the tube lens (L5) until the projected pattern was again in focus on the fluorescent coverslip and on the secondary CCD camera (**Fig**. **S2a**, *Right*). Using this strategy, we iteratively constructed an empirical calibration curve that relates the diffuser displacement to the position of the photo-stimulation plane in the sample with respect to the focal/image plane of the principal objective (**Fig. 1k**). As we moved the photo-stimulation pattern away from the focal plane of the objective, we measured a slight increase in the lateral (<10%) and axial (<20%, **Fig. S2c**), which we further accounted for by applying corrections in the control software. Over the axial displacement interval sampled, we report a linear relationship between translating the objective axially and the compensatory displacement of the diffuser required to bring the pattern back into focus, proportional to the square of the effective optical magnification of the setup.

### Relating movement of the objective to the adjusting displacement of the diffuser

Within the thin lens approximation framework, let *u* be the distance between the objective (O) and the image of the projected light pattern, *s_i_+s_o_* the distance between the objective and the tube lens (L5), *v* the distance from the diffuser to the tube lens, and *f_1_* and *f_2_* the focal lengths of the objective and the tube lens.

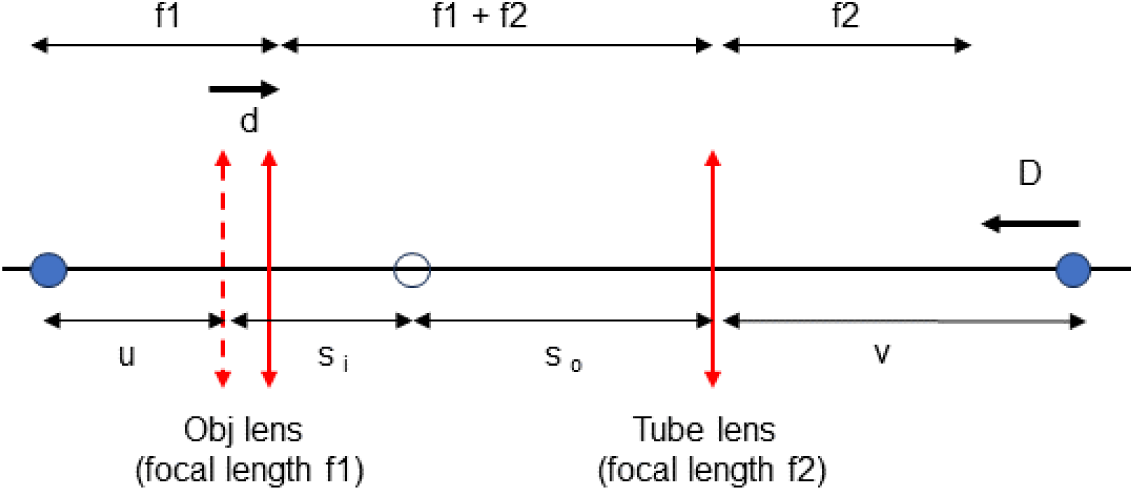

How does the displacement of the objective (*d*) relate to the displacement of the diffuser (*D*) needed to bring the projected pattern in focus in the desired z-plane in the sample, as a function of magnification (M) of the system given by the ratio of focal lengths of the tube lens and objective lens (M=f_2_/f_1_)?

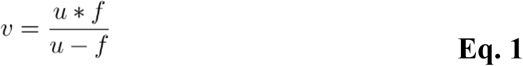

Calculating image distance (*u = f1 - d*) for the objective lens (O), when d is small:

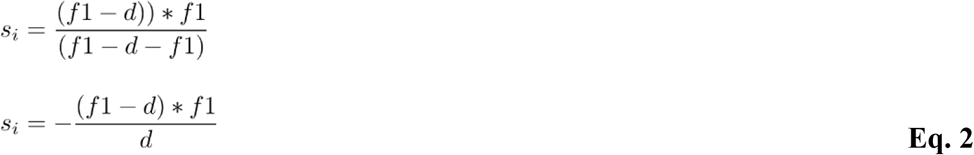

For the tube lens, the object distance is: *s_o_* = (*f*1 + *f*2 + *d*) −*s_i_*

Replacing *s_i_* from *Eq. 2*:

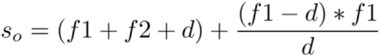

Since *d* is small, ignoring higher order terms of *d*, we obtain:

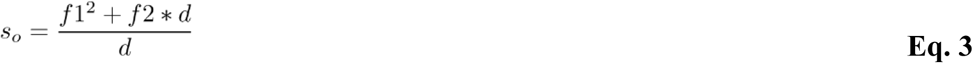

For the tube lens, image distance (*v*) is given by:

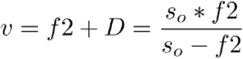

Solving for *D*:

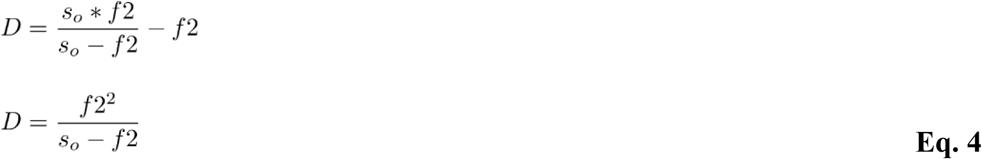

Replacing s_o_ and solving for D:

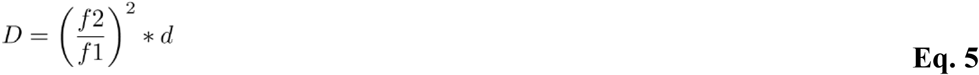

Therefore, we find that the axial displacement of the photo-stimulation plane at the sample (d) is linearly proportional to the translation of the diffuser in the optical path (D).

### Photo-stimulation and imaging calibrations

#### DMD-chip to CCD-chip to multiphoton image registration

To photo-stimulate specific locations in the sample, we determined where a given *pixel on the DMD* falls on the *specimen* using the primary camera (CCD1) as a reference. After registering pixels on the DMD-chip to CCD1 chip pixels, a point-to-point correspondence between the DMD-chip and the focal plane of the principal objective was obtained. Further, the auxiliary microscope was used to map the correspondence between pixels on the *DMD* and the *specimen* in z-planes out of the focal plane of the principal objective. To map pixels in the *two photon field of view* to the *DMD*, we also used CCD1 as intermediary reference. Within this frame of reference, we targeted ROIs that were axially (z) displaced from the principal objective focal plane, and imaged either using the primary CCD camera or the two photon microscope. Pixels of the DMD-chip were mapped onto pixels of CCD1 by projecting a series of light spots generated from known DMD locations. Routinely, we placed 20 µm sized spots in a 4 x 5 grid spanning the field of stimulation. After recording the position of all these fiduciary light spots, we computed the 3 x 3 matrix **T = DC^T^(CC^T^)^−1^**, where **C** and **D** are N x 2 matrices whose rows contain the light spot coordinates on the CCD1 and DMD chips respectively. The matrix **T**, when left-multiplied with CCD1 chip coordinates, transforms CCD1 coordinates into DMD-chip coordinates. To verify this procedure, a high-density sample with 15 µm fluorescent latex microbeads was imaged using first the widefield fluorescence path (CCD1), and subsequently the two photon microscope. Individual microbeads were matched between the acquired images manually (*cpselect* tool in the MATLAB image processing toolbox). We obtained a look-up table of two photon / CCD1 images point pairs. Further, by minimizing the sum of squared errors, we calculated the corresponding transform matrix that maps two photon image pixels onto CCD1 image pixels. The absolute position of a particular location in a two photon image is dependent on both the field of view and the resolution settings (number of pixels) of the image. Several FOV and pixel resolution settings were chosen for registration, and further used accordingly when targeting ROIs within a two photon reference image. Post-registration, we verified that the targeting of the DMD-chip generated patterns was precise. The control software reproduced the desired target light spots with high accuracy within the field of stimulation, using only the two photon image or only the CCD1 image as guide; error between projected centroids light spots vs. fluorescence microbeads was less than 5 µm, **Figs. 1j**, **S1d**.

### Correcting the field of photo-stimulation power inhomogeneity

Due to the Gaussian intensity profile of the blue laser beam illuminating the DMD chip, the power towards the borders of the field of photo-stimulation is lower than in the center. To correct for this inhomogeneity, and ensure flatness of photo-stimulation across the field, we calculated a power distribution map (PDM) across the field of stimulation (**Fig**. **S1e**), obtained by sequentially projecting 25 µm spots in an 11 x 11 grid pattern covering 121 pixels coordinates across the DMD-chip, and by linearly interpolating such as to estimate the power at the remaining positions. We projected the spot pattern on the DMD chip and placed a power sensor (Thorlabs, 50mW, S120B) under the primary objective with the pattern in focus. For simple patterns, such as spatially-localized single spots, the laser power can be adjusted accordingly using the PDM look-up table to compensate for power loss at the borders. For more complex light patterns, where different regions of the pattern have different local light intensities, pulse-width modulation can be used to control the average light intensity experienced by each spot independently.

### Off-focal plane aberration correction

We often presented a spatial photo-stimulation pattern based on glomerular shapes on the brain surface, while imaging deeper structures, such as mitral and tufted cells, which resulted in small, but significant changes in the size of the projected pattern (i.e. DMD-generated optogenetic stimulation pattern is in focus above the focal plane of the principal imaging objective). We compensated for this change in magnification by another linear transformation that we measured using the auxiliary microscope mounted beneath the principal microscope. The same procedure described above for mapping DMD-chip pixels to the primary camera (CCD1) was repeated using the secondary camera (CCD2) at several focal plane displacements in steps of 100 µm (e.g. 0, 100, 200…500 µm) and a corresponding set of transformations T(0),…, T(N) from the DMD-chip pixels to the secondary camera (CCD2) pixels was obtained. For each such transformation, we isolated the effect of changing displacement by multiplying the inverse of the zero-axial displacement transformation to obtain a set of “depth transformations” B(0),…, B(N). The depth transformation for arbitrary depths was calculated by linearly interpolating between the measured transformations.

### Dichroic mirror offset

When registering the primary camera (CCD1), the dichroic mirror (DM2) must be positioned to direct light from the sample onto the primary camera. However, light from the DMD-chip is refracted as it passes through the dichroic mirror, causing the photo-stimulation pattern to be translated when DM2 is rotated by 90 degrees towards the PMT. We measured the displacement by projecting a square light pattern and recording an image with the auxiliary microscope with the dichroic positioned towards the CCD1 or the PMT. Identifying a corner of the projected square light pattern across the two images allowed using the secondary camera CCD2-to-DMD-chip registration to determine the dichroic induced offset in projection.

### Lateral spot size measurement

To calculate the full-width at half maximum (FWHM) in the lateral direction (**Figs. 1g-i**), we projected a circular spot onto a rhodamine B-coated coverslip and acquired images with the auxiliary microscope. We thresholded each image and found the center of the spot by computing a weighted centroid of the thresholded image:

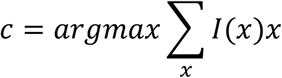

where *x* is a vector containing the pixel coordinates and *I(x)* is the brightness of the image at point *x*. The threshold was chosen heuristically by visual inspection of a single image and finding a value which isolated the light spot, and which, when changed by small amounts, did not significantly change the position of the centroid. We generated and averaged a set of line profiles passing through the centroid rotated in 1° increments over 360°. The resulting symmetric line profile was further used to compute the FWHM.

### Axial spot size measurement

To measure the axial (z) profile of the projected light spots, we brought different axial sections of the spots into the focal plane of CCD2 by stepping the principal objective up and down over an interval of ± 100 µm (for spots of diameter ≤ 40 µm) or over ± 200 µm range (for spots of diameter > 40 µm) in z increments of 1 and 2 µm respectively. We defined a region of interest (ROI) for analysis by taking all points within the FWHM of the spot in the central section. The axial profile was defined as the intensity within the ROI across all sections (**Figs. 1g-i**). To estimate the effect of spherical and other types of aberrations, we measured the axial profile of our projected spots for photo-stimulation plane z offsets of Δz = 0 – 400 µm with respect to the focal plane of the principal objective (**Fig**. **S2c**). For these measurements, we used spot diameters of 30, 60 and 90 µm respectively. Since in these configurations the diffuser is not positioned at a plane conjugate to the focal plane of the principal objective, any relative movement of the principal objective with respect to the tube lens L5 results in a slight change in magnification. The projected light spot size was corrected accordingly for the predicted increase in magnification.

### Optical alignment

Alignment of the two-photon microscope followed the typical procedure^112^. The photo-stimulation beam was further aligned on the DMD chip. In our implementation, the DMD chip was mounted onto an articulating base-ball stage (SL20, Thorlabs) that can be raised and lowered. We used a periscope (RS99, Thorlabs) formed by two mirrors placed on kinematic mounts to center the photo-stimulation laser beam on the DMD chip. By applying an all-white pattern to the DMD-chip, a coarse alignment was obtained with the light reflected off the DMD-chip into the microscope and the photo-stimulation beam centered on the chip. One point of potential difficulty in the alignment procedure is given by the diffractive behavior of the DMD-chip when illuminated with coherent light. The face of each individual micro-mirror is offset with respect to the plane of the entire array of mirrors. Because of this, the DMD-chip behaves like a blazed diffraction grating. We aligned the blue laser beam such that most of the power is sent into a particular diffraction order. For a one-dimensional grating, the angle *θ_m_* of diffraction order *m* is given by:

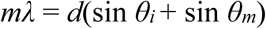

while *θ_r_*, the angle at which the reflected laser power leaves the DMD chip is given by:

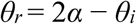

where *θ_i_* is the incident angle with respect to the normal of the grating, *λ* is the wavelength, *d* is the grating spacing, and *α* is the offset or “blaze angle” of the mirror face relative to the grating. Since the micro-mirrors rotate along one diagonal of the mirror, the projection of the correct incident angle onto the mirror face must lie orthogonal to this diagonal. This line was determined by toggling all the mirrors between the on and off positions (setting an all-white or all-black image) and observing where the light is redirected.

To make final adjustments to the position, we compared the power in the brightest diffraction order to the four adjacent ones. We then adjusted the incident angle of the beam by tilting or translating the DMD-chip mount and compensating with the periscope mirrors to keep the photo-stimulation beam centered until the power in the brightest spot was maximally bright relatively to its neighbors. After optimizing the incident angle, the brightest spot was ∼ 60% as powerful as before the DMD chip. A circular iris (SM2D25, Thorlabs) was used to block out light from other diffraction orders. Applying a cross-shaped or small circular pattern to the DMD chip is useful for determining the center of the field of stimulation. Finally, the dichroic mirror (DM1, **Fig. 1b**) was used to align the center of the photo-stimulation pattern to the same vertical line as the two-photon beam, and the primary camera (CCD1) and PMTs were placed and aligned as described above.

### Propagation of patterns through brain tissue

We used the auxiliary microscope to characterize how the lateral profile of the one photon extended (2D) photo-stimulation patterns deteriorates after propagation through brain slices of different thicknesses, following the procedure described in *Papagiakoumou et al., 2013*^113^. After focusing the photo-stimulation pattern on the fluorescent coverslip so that it was in focus on both the primary and the secondary cameras, we placed a slice on the top of the coverslip with the pattern illuminating the olfactory bulb. The photo-stimulation light travelled through the slice and was scattered by the tissue before the pattern was imaged on the coverslip and the secondary camera (CCD2). We took a total of six measurements for each slice thickness (three positions, two slices). We used as reference for normalization the configuration when no slice was present (0 thickness) and defined the ROI for analysis as the set of points within the FWHM when no slice was present. We then calculated the light intensity and the standard deviation of pixel values within the ROI and compared it to values obtained for different slice thicknesses (**Figs**. **S3a-e**). Patterns remained distinct in slices up to ∼150 µm thick (**Figs**. **S3a-e**), albeit substantial fluorescence loss (**Fig**. **S3e**). We note that using the diffuser, besides confining the axial spread of photo-stimulation, also increases the homogeneity of the projected light patterns in scattering media (**Figs**. **S3a-e**). Without the diffuser in the optical path, patterns appeared speckled: standard deviation of pixel intensities within the pattern was higher compared to same thickness samples with the diffuser in place (**Figs**. **S3a-e**). This effect can be explained intuitively by taking a ray-tracing point of view on image formation. With the diffuser in the optical path, due to angular dispersion, light emitted from a single point in the pattern travels along more paths through the tissue compared to the no-diffuser condition to reach at a given camera pixel (**Fig**. **S3c**). Using the diffuser increases homogeneity in light intensity across the field of stimulation as this configuration is less sensitive to variations in scattering from local structure in the tissue. The axial profile of the projected spots substantially deteriorated with increasing in slice thickness beyond 100 µm (**Fig**. **S3f**). As such light scattering constrains using this approach for pattern photo-stimulation of superficial structures, within 100 µm from pia, while imaging ensuing responses in deeper layers.

### Phosphorescence artifacts

We observed a small slowly-decaying PMT signal upon illumination with the blue photo-stimulation laser in the absence of a fluorescence specimen. By progressively blocking off different parts of the optical path, we determined that the main contributors to this signal were the objective and tube lens (O and L5 in **Fig. 1**). Upon replacing the N-BK7 glass tube lens (LA1384-A) with a fused silica lens (LA4545-A) with the same focal length, the signal dropped substantially (**Fig**. **S1e**). The artifact was not affected by the type of anti-reflective coating used (**Fig**. **S1e**). We tested several objectives (Olympus 20x 1.00 NA XLUMPlanFLN, Olympus 20x 0.95 XLUMPlanFL, Olympus 25x 1.05NA XLPlanN) and determined that they contributed differentially to this signal. Of these, the 25x fared best followed by the 1.0 NA 20x and the 0.95 NA 20x (data not shown). Following this optimization, when comparing identified cells to background areas in our imaging experiments, we could not detect any spurious signals in background selected areas.

### Mice

For the *in vivo* experiments described here, we included data from 12 OMP-Cre x ReChr x Thy1GCaMP6s mice, 3 DAT-Cre mice and 4 DAT-Cre x Thy1-GCaMP6f mice. For experiments to map mitral/tufted cell parent glomeruli and identify *sister* cell cohorts, triple mouse crosses (OMP-Cre x Rosa26 CAG-LSL-ReaChR-mCitrine^86^ x Thy1GCaMP6s_4.3^88^) were used to express ReaChR-mCitrine and GCaMP6f/s in the OSNs and respectively mitral and tufted cells (**Figs**. **2**,**3, S4**, **S5**). For experiments expressing ChR2 and GCaMP6f in DAT+ cells (**Figs. 4a-c**, **S6a-b**), we used mice expressing Cre recombinase in DAT+ neurons and approximately 300 nl of a mixture of AAV (serotype 2.9, UNC Vector Core) virus carrying DIO-ChR2-mCherry and DIO-GCaMP6f was injected into the olfactory bulb at a depth of 300 µm as previously described^41^. For glomerular stimulation, for each light stimulus other than control stimuli, we fit a function of the form *f(d) = Ae^−d/b^*where *f* represents the light-evoked fluorescence change in a region of interest and *d* is the distance from that region to the center of the projected light mask. The width constant *b* of this exponential fit gives a measure of the spread of activity evoked by that stimulus. For each light intensity sampled, we repeated the analysis in two different fields of view, and the relationship between activation width and light intensity is shown in **Figs. 4c**, **S6e**. As in the case of OSN terminals photo-stimulation experiments, we flashed single 100 ms sub-glomerular light masks (20 µm diameter) between the imaging frames and recorded calcium transients in the DAT+ cells via multiphoton imaging. For analysis, the field of view was divided into a 20 by 20 grid (15 x 15 µm grid units), and the response was defined as the average change in fluorescence in each unit of the grid during the two frames (10 Hz) immediately following the light stimulus. We tested a range of photo-stimulation light intensities ranging from 0.5 to 10.0 mW/mm^2^. We tuned up the intensity of the fluorescence response, while aiming to maintain spatial specificity, as indicated by the photo-activated area remaining largely confined to the chosen glomerulus. We quantified the spread of activity by characterizing the relationship between the DAT+ cell response strength and the distance from the centroid of the light stimulus, which was well-fit by exponential decay. Within the range of light intensity used, the characteristic width constant varied from 25 µm for the lowest to 62 µm for the highest.

For experiments investigating the impact on mitral/tufted cell activity of glomerular GABAergic/DAT+-neurons (superficial short axon cells), DAT-Cre mice were crossed to Thy1GCaMP6f_5.11^88^ mice (**Figs**. **4**,**5 S6-8**). DAT-Cre x Thy1-GCaMP6f mice were retro-orbitally injected^114^ with 5 × 10^12^ vp of AAV-PHP.eB:Syn-FLEX-rc[ChrimsonR-tdtomato], which was prepared as previously described^100,115^. Viral particles were diluted in saline to achieve a final volume of 100 μL per injected mouse. All animal procedures conformed to NIH guidelines and were approved by the Animal Care and Use Committee of Cold Spring Harbor Laboratory.

### Surgical procedures

Mice were injected with NSAID Meloxicam 0.5 mg/Kg (Metacam®, Boehringer Ingelheim. Ingelheim, Germany) 24 hours prior the surgical procedure, at the onset of surgery, and for 2 days *post* each surgical procedure. Depending on the recovery progression, the NSAID treatment was maintained until mice showed alert, and responsive behavior. Before each stereotaxic surgery, mice were anesthetized with 10% v/v Isofluorane (Cat# 029405. Covetrus. Portland, ME, US). For the chronic window and headbar implantation procedures, mice were anesthetized with a ketamine/xylazine (125 mg/Kg - 12.5 mg/Kg) cocktail. During surgery, the animal’s eyes were protected with an ophthalmic ointment (Puralube®. Dechra. Nortwich, England, UK). Temperature was maintained at 37 °C using a heating pad (FST TR-200, Fine Science Tools. Foster City, CA, USA). Respiratory rate and lack of pain reflexes were monitored throughout the procedure. Chronic window implant surgeries were supplemented with dexamethasone (4 mg/Kg) to prevent swelling, enrofloxacin (5 mg/Kg) to prevent bacterial infection, and carprofen (5 mg/Kg) to reduce inflammation.

### Chronic implantations

After recovery from the stereotaxic surgery, mice were implanted with a custom titanium head bar attached with C&B Metabond Quick adhesive luting cement (Cat# S380. Parkell. Edgewood, NY, USA), followed by black Ortho-JetTM dental acrylic application (Cat# 1520BLK. Lang. Chicago, IL, USA) and with a cranial window on top of the olfactory bulb as previously described ^116,117^. Special care was taken to remove the bone under the inter-frontal suture and remove small bone pieces at the edges. During surgery, the exposed olfactory bulb was continuously protected and cleaned of blood excess using artificial cerebrospinal fluid (aCSF) and aCSF-soaked gelfoam. Once both hemibulbs were exposed and clean, a fresh drop of aCSF was placed on top, followed by a 3 mm round cover glass, which was gently pushed onto the OB surface to minimize motion artifacts in further experiments. Once in place, the coverslip was sealed along the edges with a combination of VitrebondTM, Crazy-GlueTM, and dental acrylic to cover the exposed skull. Mice recovered for ∼ 7 days before imaging experiments.

### Odor delivery

Odors were presented using a custom-built odor machine as described elsewhere^92,118^. Undiluted or oil diluted (1:10 or 1:100) odorants were placed in glass vials and airflow through the vials controlled by solenoid valves. The output odor stream was further mixed with clean air in a 50:1 ratio, then this diluted stream was further mixed with clean air at a 5:1 ratio before being sent through short teflon coated tubing placed in front of the mouse’s snout. The 12 vials in the machine were split into three banks of four vials. Each bank could be independently switched between sending odorized air and clean air to the mouse, in principle allowing combinations of up to three odors. For assessing the effect of modulating GABA-ergic/DAT+ interneurons onto mitral and tufted cells responses, we used the following odors: isoamyl acetate, acetaldehyde, valeraldehyde, allyl tiglate, ethyl heptanoate, heptanal, hexanol, 2-hexanone, ethyl tiglate, 1,4 cineole, acetophenone.

### Data pre-processing

#### Movement correction and ROI selection

Rigid registration in Fiji/ImageJ (template matching) was applied to the acquired fluorescence time-lapse stacks. Images were visually inspected to select a motion-free sequence of frames and create a mean reference image to which we registered each image stack corresponding to a given trial. ROI selection was performed manually (ImageJ): we used both the average and standard deviation projections of the registered images to draw ROIs around the glomeruli and respectively mitral and tufted cell bodies in the FOV.

#### Bleaching correction

Post ROI extraction (glomeruli, mitral/tufted cells), we applied fluorescence bleaching correction, assuming that: (1) bleaching follows a first order exponential decay across trials for each individual ROI; (2) during the baseline period, any measured activity is random. Given these assumptions, by averaging the fluorescence traces across trials of an ROI, the only robust trend potentially present in the baseline is the characteristic bleaching for that ROI. We checked that the baseline fluorescence of a given ROI is comparable across trials within a session. The procedure used for each ROI was as follows: (1) we fitted an exponential decay function to the fluorescence traces of individual trials, (2) subtracted the fitted function from the ROI trace, (3) averaged the fluorescence traces across all trials for that ROI (aligned to stimulus presentation).

#### Normalization

For each ROI and trial, we normalized the traces individually, using a fluorescence baseline period (average of ten frames just before the stimulus onset as baseline fluorescence, F0). For each time point, fractional changes over the baseline fluorescence for the average trace were calculated as dF/F0.

#### ROI responsiveness and classification

On average, stimuli were presented across 4-5 repeats when imaging glomerular responses to optogenetic stimulation, and across 6-10 repeats when sampling the responses of mitral and tufted cells. To evaluate the responsiveness of each glomerulus, we obtained a distribution of average dF/F0 values from baseline periods, and used 99 percentile value of the distribution as response (significant signal) threshold. The baseline reference distribution was obtained from averaging dF/F0 values quantified over 10-frames intervals extracted from the baseline period and accumulated across all trials. For each ROI, we also obtained peak dF/F0 values during the optogenetic stimulation frames (1-2 frames). For each ROI, we compared the average response across trials with the baseline reference distribution. If this value crossed the 99^th^ percentile of the baseline distribution, the ROI was classified as responsive (*enhanced*).

To evaluate the responsiveness of each mitral and tufted cell, we searched for the peak of a monotonous response (signal) within 5 frames following optogenetic stimulation onset. Similarly, we obtained a distribution of average dF/F0 values from baseline periods using 4 frames just before the stimulation and accumulating across all trials. We compared the peak response value to the standard deviation (SD) of the baseline. For calling *enhanced* responses, we used 4 SD of the baseline as response (significant signal) threshold; for calling *suppressed* responses, we used 3 SD of the baseline as response threshold to account for the intrinsic lower range of negative deflections from baseline. Cells that had slow ramping and prolonged negative responses (response slope <15° with respect to baseline) were not included in further analyses. To assess response reproducibility across trials, for each optogenetic stimulation mask, for each trial, we built a *trial response vector* (using 5-frames from the stimulus onset) and computed the Pearson correlation with respect to the response vectors of the other trials. If for a given trial the average correlation with respect to the response vectors of the other trials, the trial was rejected. For an ROI to be considered for further analysis, at least 2 repeats had to pass the criterion. Cells that had slow responses, which occurred only after offset of light stimulation were not included for further analysis to minimize the contribution of disinhibitory effects across the bulb network.

#### Self-response correlations

To obtain a measure of reliability for ensemble mitral cell responses across different trials to the same DAT+ glomerular photo-stimulation mask, and respectively to the same odorant, using bootstrapping (50 times) in each iteration split the trials into two random subsets of trials, and calculated ‘self’ response correlations, whose average values were 0.81 ± 0.14 for light masks and 0.87 ± 0.10 for odor responses of the same MC ensembles (**Fig. 4h**).

For each mitral and tufted cells, the *odor responses* (**Fig. 5**) were calculated as described above using 10-frames periods to build a distribution of baseline values and accumulating across trials. We also used 10-frames periods starting from stimulus onset to search for the response peak. To calculate the *light modulation of the odor response* to DAT+ glomerular photo-stimulation masks (**Figs. 4e-h**, **S8**), we used as reference the frame just before light stimulation onset. We asked whether in the 2 frames following light onset, we could identify a significant increase/dip in signal compared to this reference dF/F value. A change in signal (defined delta between reference and signal during light stimulation) was considered significant if it amounted to more than 10% of the average peak odor response amplitude (preceding light onset), and was also larger than 3 SDs of the baseline fluorescence (preceding odor onset). To assess response reproducibility across trials, for each stimulus, we performed correlation analysis across trials as described above.

#### Assessing glomerular response spatial extent and specificity

In **Fig**. **S4c** *i* (Right), for each photo-stimulation mask, we normalized the sum total area of the significantly responsive glomerular pixels. We multiplied the sum total area of these responsive pixels by the ratio between the area of the photo-stimulation mask and the surface area of target glomerulus surface. We defined the anatomical outline of each glomerulus in the field of view manually using its baseline fluorescence. To determine the specificity of glomerular responses across light intensities we computed two metrics. First (blue trace, **Fig. 2f**), we normalized responsive pixels within the boundaries of the target glomerulus by all responsive pixels located within the boundaries of any glomeruli across the FOV. Second (red trace, **Fig. 2f**), we normalized the significantly responsive pixels within the target glomerulus by the sum area of all responsive pixels within the FOV, including those spanning the juxtaglomerular space, surrounding the glomeruli. The specificity index was high when only glomerular pixels are considered (**Fig. 2h**), yet 5-8 % of significant pixels were in off-target glomeruli, potentially due to spurious stimulation of fibers of passage.

#### Correlation analysis and shuffled controls

For each pairwise correlation analysis (**Figs**. **4h**, **5f-h**, **S7b, S8b**), we ran 15 repetitions of the shuffling controls and compared the experimental data with the shuffled data (Wilcoxon signed-rank test).

#### Statistical tests

Depending on the properties of the analyzed data, different statistical tests were used to evaluate the differences between data groups. The information for each statistical comparison is detailed in the legends of each figure panel. ‘Two-sample unpaired Student’s t-test’ was used to evaluate significant differences in the means of two normally distributed data sets. ‘The normality of the data distribution was checked using a ‘One-sample Kolmogorov-Smirnov test’ for each data group before the testing differences with the ‘Student’s t-test’. A two-tailed 95% confidence interval was used for both Student’s t-test. Significant differences between two non-normally distributed data sets were evaluated with a nonparametric ‘Wilcoxon signed-rank test.’ The linear correlation between two data sets was assessed by computing the Pearson correlation coefficient. When needed, the coefficient of determination (R^2^) was used to evaluate how well the compared data sets fit a linear regression model. All the statistical comparisons were tested using MATLAB® using the following functions for each test: ‘Two-sample unpaired Student’s t-test’: ttest; ‘One-sample Kolmogorov-Smirnov test’: kstest; ‘Multiple comparisons of means test’: multcompare; ‘Two-sample Kolmogorov-Smirnov test’: kstest2; ‘Pearson correlation coefficient’: corrcoef. ‘Wilcoxon signed-Rank’: signrank.

