## Supplementary Material for "Axially decoupled photo-stimulation and two photon readout (*ADePT*) for mapping functional connectivity of neural circuits"

#### Supplementary Figure 1

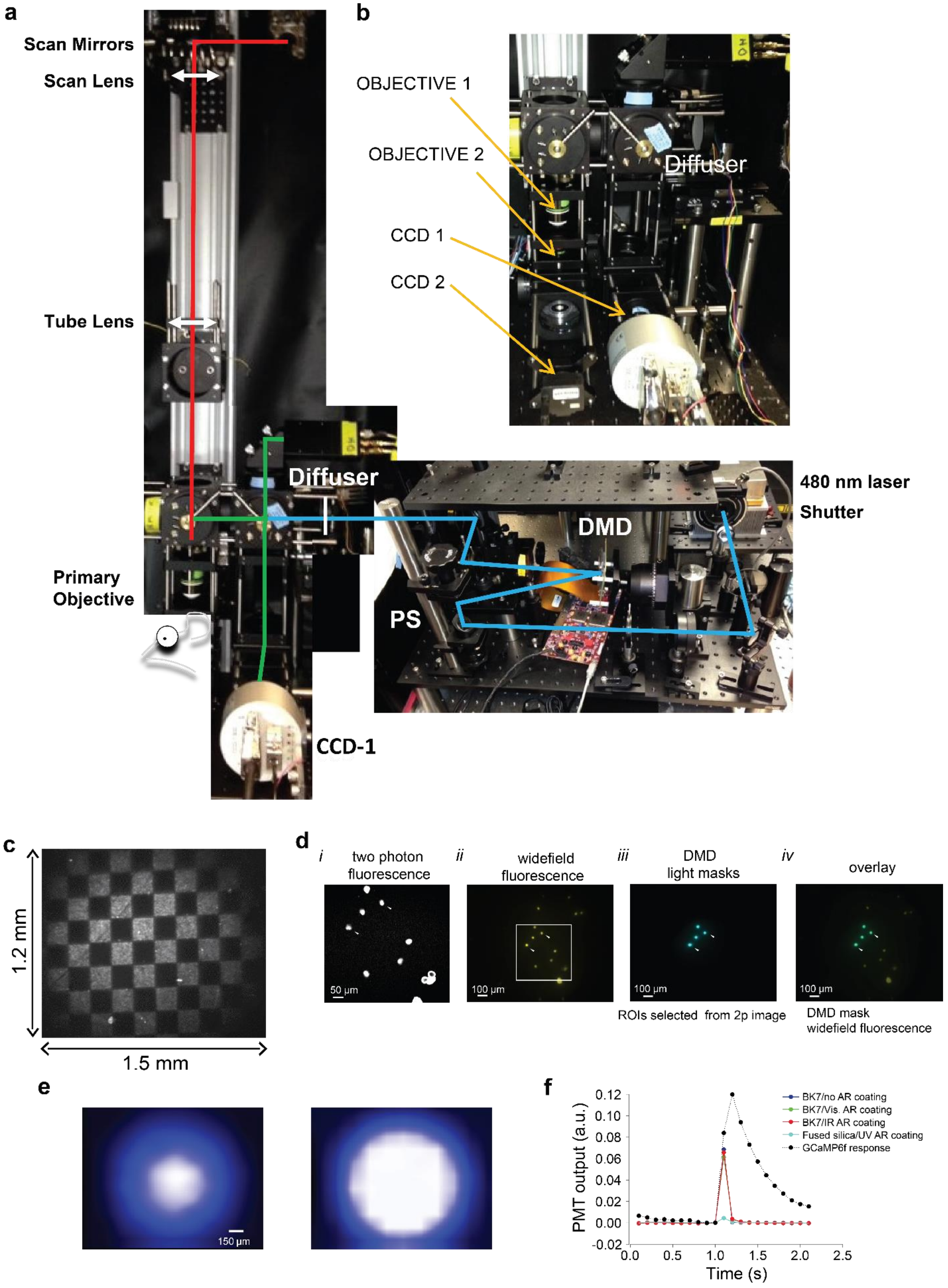

***Figure S1. ADePT photo-stimulation rig, mapping DMD-to-CCD-to-2p images, and calibration of the DMD-modulated photo-stimulation patterns**. Corresponding to **Fig**. **1**. **(a,b)** Photo-stimulation and imaging rig pictures: CCD1/CCD2 – primary and secondary CCD cameras; DM, dichroic mirrors; DMD, digital micro-mirror device; I, iris diaphragm; L1-L12, lenses; PMT, photomultiplier tube; PS, periscope. Photo-stimulation (2D-light mask from a digital micro-mirror device illuminated with a solid-state CW 480 nm laser) and two photon imaging are confined to different optical z-planes that can be flexibly and independently adjusted by translating the diffuser and respectively the primary objective. **(c)** Fluorescence image of DMD-modulated light checkerboard projected on a 1 µm thin rhodamine B coated slide across the 1.2 x 1.5 mm field of stimulation (FOS) acquired using the secondary (inverted) microscope (CCD2). **(d)** DMD chip-to-CCD camera-to-2p microscope registration. (i) two photon fluorescence image of pollen grains; (ii) widefield fluorescence image (full field illumination) of a larger field of view including the pollen grains from **(i)** marked by the square; (iii) ROIs selected from the two photon image were used to generate light masks on the DMD-chip that were further projected at the focal plane of the principal objective and imaged using the primary CCD camera (CCD 1); (iv) overlay of the DMD-chip generated photo-stimulation masks and widefield fluorescence image of pollen grains; fiduciary arrows mark indicate the location of same two pollen grains in the widefield fluorescence and two photon images. **(e)** Applying a correction mask for light power inhomogeneities arising due to the Gaussian profile of the photo-stimulation laser beam and aberrations along the optical path. A power distribution map (PDM) across the field of stimulation was obtained by sequentially projecting 25 µm spots in an 11 x 11 grid pattern covering 121-pixel coordinates across the DMD-chip, and by linearly interpolating such as to estimate the power at the remaining positions. A power meter (Thorlabs) 50 mW sensor was placed in the focal plane of the primary objective with the projected DMD pattern in focus (Methods; Left/Right: uncorrected vs. corrected images). **(f)** The PMT output displayed a slowly-decaying (phosphorescence) signal upon illuminating a cover glass placed in the focal plane of the primary objective with the blue laser. By progressively blocking off different parts of the optical path, we determined that the main contributors to this signal were the objective and tube lens (Obj and L5 in **Fig**. **1**). Replacing an N-BK7 glass tube lens with a fused silica lens with the same focal length, the signal dropped substantially. The artifact was not affected by the type of anti-reflective coating used.*

#### Supplementary Figure 2

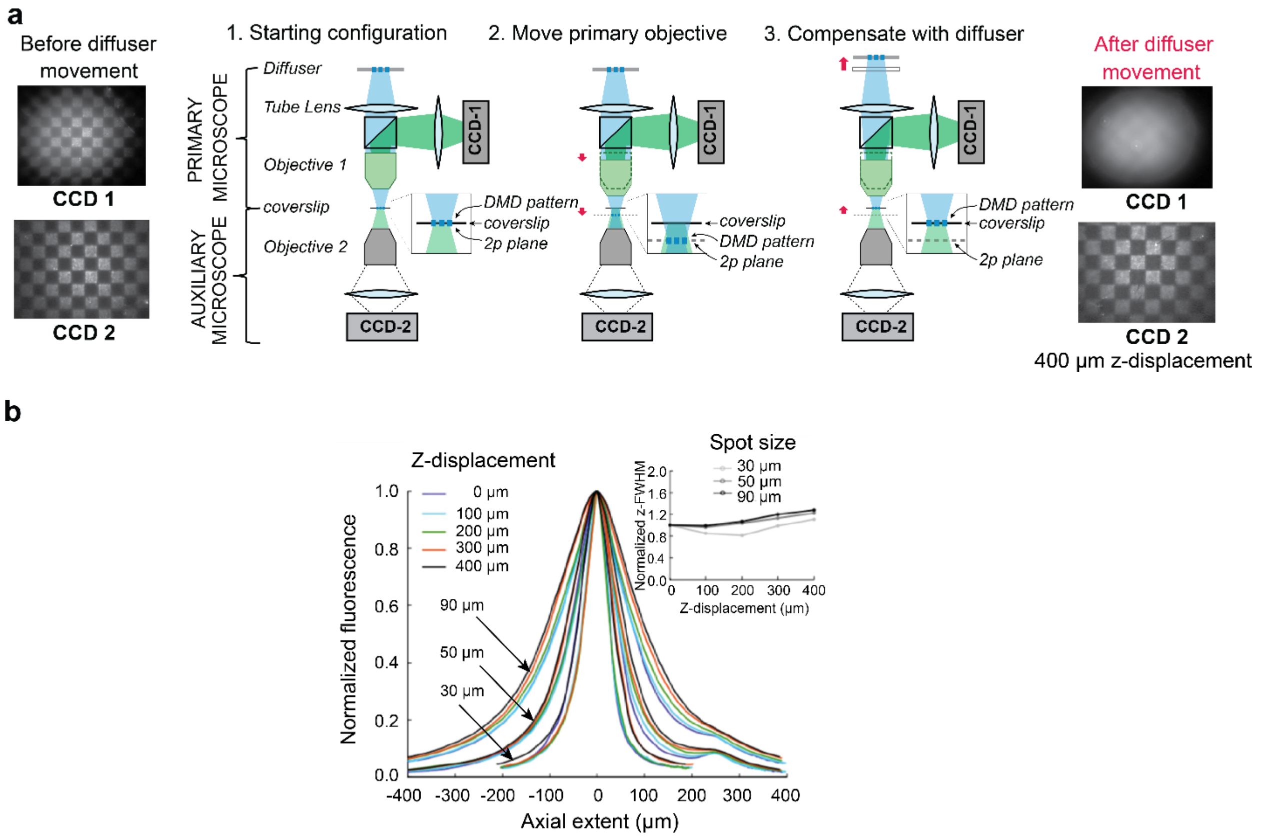


***Figure S2. Decoupling the photo-stimulation and imaging z-planes using a translatable diffuser****. Corresponding to* ***Fig****. 1.* ***(a)*** *Schematic cartoon describing using of the primary and secondary (auxiliary) microscopes to calibrate the decoupling of the photo-stimulation and optical-readout planes. A thin (1 µm) rhodamine B fluorescent coverslip sample is placed in the focal plane of both primary and secondary microscope objectives, where a checker-board pattern is projected using the DMD-chip (Step 1). Lowering the primary objective by a given amount translates the pattern axially below the focal plane of the secondary objective (which remained stationary), and produces a blurred image of the checker-board pattern on the secondary (CCD2) camera (Step 2). The projected light pattern is brought back into focus on CCD2 by compensating with translating the diffuser along the optical path (i.e. moving the diffuser away from the back-aperture of the primary objective, which defocuses the image on the primary (CCD1) camera (Step 3).* ***(b)*** *Normalized axial light intensity profile of spots. Measurements for spots of three different diameters (30, 50 and 90 µm), indicated by the three clusters of color traces. Each line color represents a different diffuser offset, expressed as degree of axial displacement between the photo-stimulation and focal plane of the primary objective in 100 µm steps (0 to 400 µm). (Inset) Measured change in axial (z) FWHM of the light profiles in* ***b.*** *as a function of axial displacement between photo-stimulation and imaging planes relative to no-offset (z-overlapping) condition, which we account for by applying corrections in the control software.*

#### Supplementary Figure 3

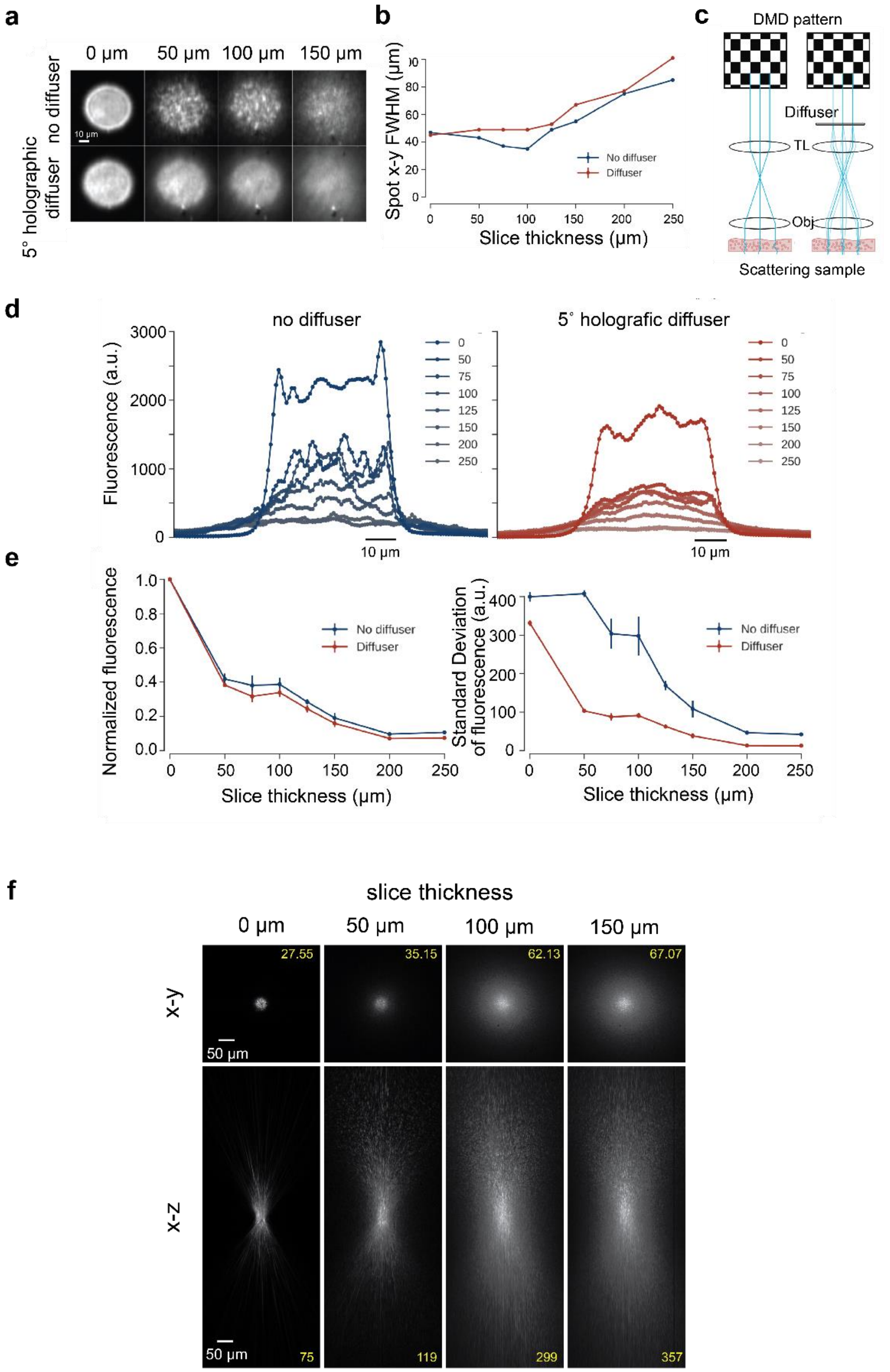

***Figure S3. Lateral and axial changes of projected patterns as a function of tissue depth penetrance of ADePT photo-stimulation***. *Corresponding to* ***Fig. 1***. ***(a)*** *50 µm circular spot projected onto a rhodamine B coated coverslip through acute mouse brain slices of different thickness, from 0–150 µm. Note the difference in homogeneity of the light patterns in the presence/absence of the diffuser.* ***(b)*** *Dispersion of the pattern assessed by calculating the lateral (x-y) FWHM increases with slice thickness, but is comparable in the presence (red) and absence the diffuser (blue).* ***(c)*** *Cartoon schematics depicting (ray diagram view) differences in pattern homogeneity through scattering tissue; in the absence of the diffuser, the example rays (drawn parallel to the optical axis) from three points within the pattern travel straight paths before propagating and scattering through the tissue; the diffuser diverges the example rays across multiple optical paths (spanning a solid angle cone) which further propagate through the tissue across a wider surface area, reducing the effect of variation in scattering properties along a particular path in the final image.* ***(d)*** *Lateral extent line profiles through the center of the spot for different slice thicknesses without (Left) and with (Right) the 5° holographic diffuser.* ***(e)*** *(Left) Normalized light intensity of 50 µm circular spots decreased at comparable rates with and without the diffuser with increasing slice thickness. Measured values were normalized to signal obtained in the absence of the slice (0); (Right) Standard deviation (SD) of fluorescence within the spots as a function of slice thickness (0–250 µm). Images taken with the diffuser in the photo-stimulation path have less fluctuation in brightness (fewer speckles).* ***(f)*** *Changes in lateral (x-y) and axial (x-z) FWHM of a 30 µm diameter circular spot projected on the fluorescent coated coverslip through acute brain slices (0-150 µm). Same optical configuration without a diffuser results in a light pencil (****Fig. 1g****). Patterned photo-stimulation is constrained by light scattering to structures within 100 µm from surface.*

#### Supplementary Figure 4

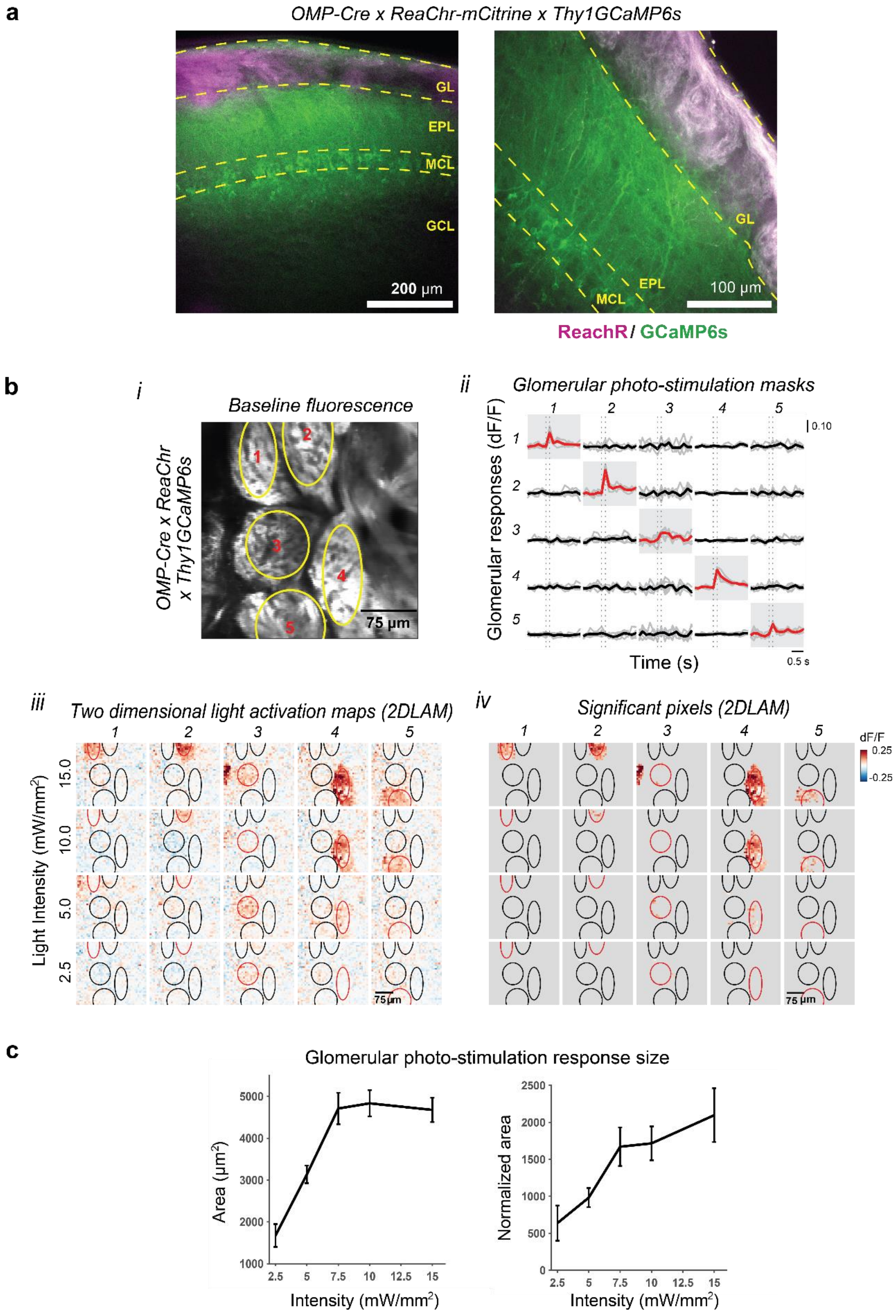

***Figure S4. Targeting individual glomeruli by photo-stimulation of OSN terminals and monitoring glomerular responses of sister mitral and tufted cell dendrites.*** *Corresponding to* ***Fig. 2***. ***(a)*** *Confocal image of olfactory bulb sagittal slices showing expression of ReaChR-mCitrine in the OSNs (glomerular layer) and GcaMP6s in the mitral and tufted cells in OMP-Cre x ReaChR-Citrine x Thy1GCaMP6s.DKim4.3 mice.* ***(b)*** *(i) Example baseline fluorescence in a glomerular field of view (FOV) from an OMP-Cre x ReaChR-citrine x Thy1-GCaMP6s mouse; (ii) Response matrix of fluorescence changes (individual repeats in gray, and mean signal as thicker lines) corresponding to five glomerular regions of interest versus the five (matching) glomerular light masks (i) projected on the OB surface. Red traces mark average responses of the target glomerulus; (iii) Fluorescence changes (two dimensional glomerular light activation maps, 2DLAMs) evoked by photo-stimulating the five targeted glomeruli (i) across a range of four light intensities (2.5, 5.0 and 10.0 and 15.0 mW/mm$^2$; 2 x 150 ms stimulus duration/frame); (iv)* *Statistically significant photo-stimulation responses shown as rectified 2DLAMs. Non-significant responses were thresholded to blue (Methods). All five glomeruli targeted for photo-stimulation were responsive.* ***(c)*** *Average lateral extent of glomerular fluorescence response while varying the light intensity (2.5 to 15 mW/mm$^2$). (Left) Surface area of significantly responsive pixels; expected surface area, assuming average glomerulus diameter of 75 µm and no activation of cells surrounding the glomerulus, is ~4,400 µm$^2$. (Right) Responsive pixels area normalized using the ratio between the photo-stimulation mask and target glomerulus areas (Methods).*

### Supplementary Figure 5

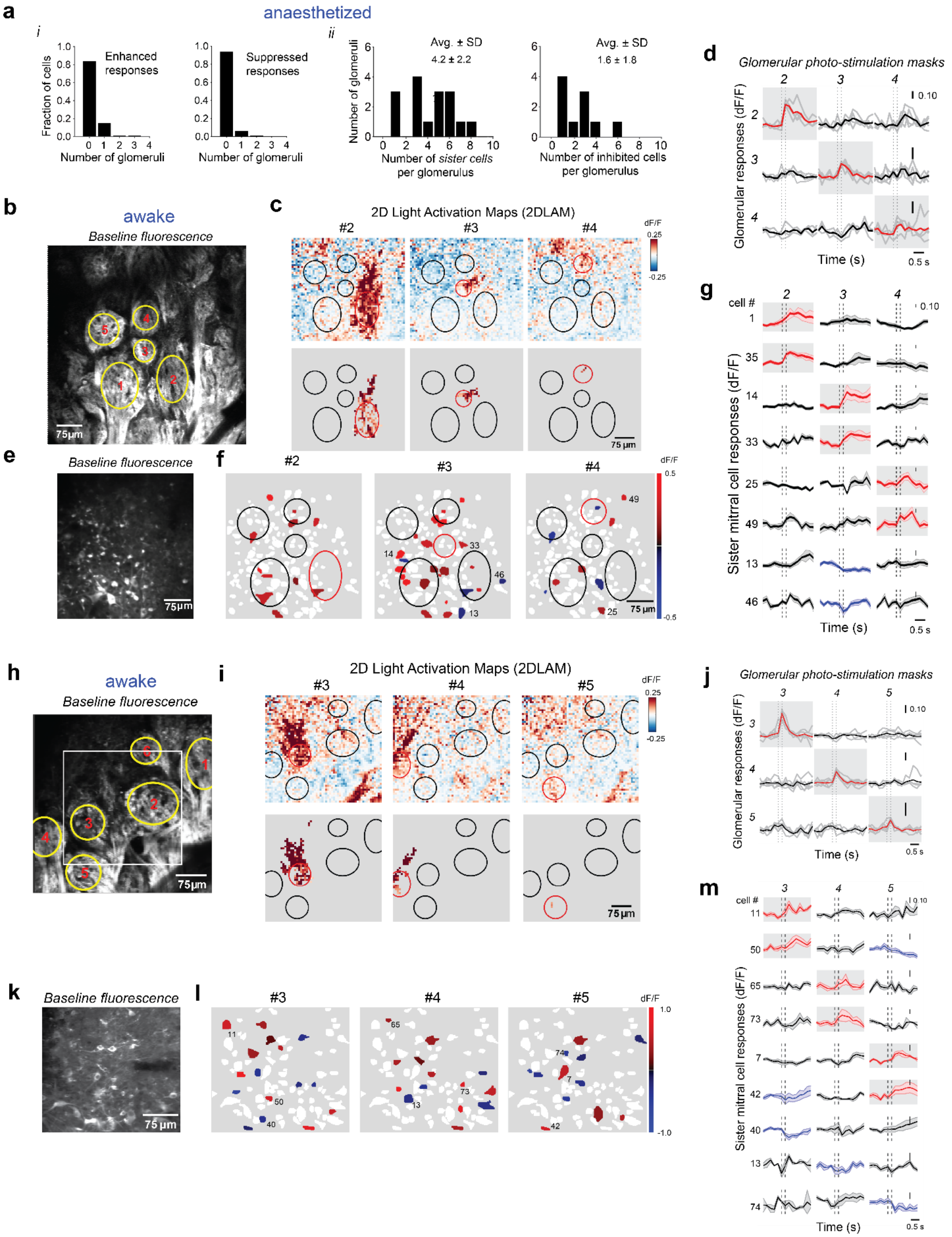

***Figure S5. Identifying sister mitral and tufted cells via axially decoupled glomerular stimulation and two photon fluorescence imaging of cell body responses**. Corresponding to **Fig. 3**. **(a)** anaesthetized, **(b-m)** awake. **(a)** (i) (Left) Specificity of enhanced responding mitral and tufted cells as a function of targeted glomerulus (N=5 mice; 16 responsive and specific glomeruli / 24 photo-stimulated target glomeruli; 357 mitral & tufted cells). Note that most cells were non-responsive, or responded specifically to the photo-stimulation of the target glomerulus (10.3 % of cells had off-target responses to more than one glomerulus; Methods); (Right) Same for suppressed responses. (ii) (Left) Number of observed sister cells (enhanced responses) per glomerulus photo-stimulation across fields of view (Left; Avg. ± SD,* 4.2 ± 2.2; N = 6 FOVs; 5 mice*); (Right) Number of cells whose baseline fluorescence was suppressed upon photo-stimulation of the same glomeruli (Right; Avg. ± SD*, 1.6 ± 1.8; N = 6 FOVs; 5 mice*). **(b)** Resting glomerular fluorescence (ReaChR-citrine and GCaMP6s), awake; **(c)** Fluorescence changes (raw, Top, and significant pixels glomerular 2DLAMs, Bottom) evoked by photo-stimulation of three target glomeruli (7.5 $mW/mm^2$; 100 ms light on/trial); **(d)** Response matrix of fluorescence change traces (individual repeats in gray, and mean signal as thicker lines) corresponding to three glomerular regions of interest (#2,#3,#4) versus the 3 (matching) glomerular light masks projected on the OB surface. Red traces mark responses of the targeted glomerulus in a given trial; stimulation of glomeruli #1 and #5 resulted in non-specific responses. **(e)** Resting fluorescence (GCaMP6s) of mitral cell bodies at the same x-y coordinates, 180 μm below the glomerular field of view shown in (**b**); **(f)** Significant cell body responses evoked by photo-stimulation of three target glomeruli (**g**,ii); (iii) Average fluorescence time traces corresponding to nine example mitral cells in each heat map, whose positions are indicated by numbers in (ii). Three example pairs of daughter sister cells (1, 35 with respect to parent glomerulus #2; 14, 33 with respect to parent glomerulus #3; and 25 and 49 with respect to parent glomerulus #4) are shown. In addition, glomerular photo-stimulation triggered spatially distributed suppressed responses (e.g. cells 13, 46). **(h-j)** Same as (**b-d**) for another awake experiment: photo-stimulation of three target glomeruli (2.5 $mW/mm^2$; 50 ms light on/trial); **(i)** Significant cell body responses evoked by photo-stimulation of three target glomeruli (**h**); **(j)** Response matrix of fluorescence change traces (individual repeats in gray, and mean signal as thicker lines) corresponding to three glomerular regions of interest (#3,#4,#5) versus the three (matching) glomerular light masks projected on the OB surface. Stimulation of glomeruli #1 and #2 resulted in non-specific responses. **(k-m)** same as (**e-g**) for sampling mitral cell bodies at the same x-y coordinates, 190 μm below the glomerular field of view shown in (**h**); **(l)** Significant cell body responses evoked by photo-stimulation of three target glomeruli (**h**); **(m)** Average fluorescence time traces corresponding to nine example mitral cells in each heat map, whose positions are indicated by numbers in **(l)**. Three example pairs of sister cells (11, 50 with respect to parent glomerulus #3; 65, 73 with respect to parent glomerulus #4; and 7 and 42 with respect to parent glomerulus #5) are shown. Glomerular photo-stimulation also triggered spatially distributed suppressed responses (e.g. cells 13, 40, 42, 50, 73, 74), denser than in anaesthetized mice.*

#### Supplementary Figure 6

**a**

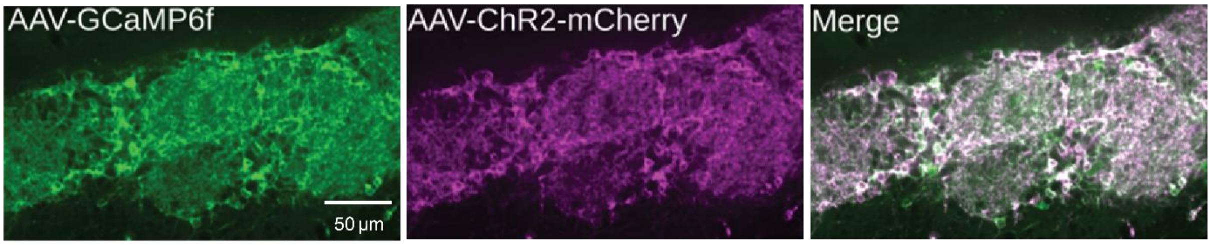


**b**

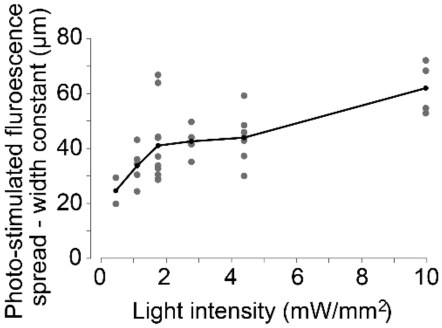


***Figure S6. Optogenetic control of DAT+ (superficial short axon) cells within individual glomeruli**. Corresponding to **Fig**. 4. **(a)** Confocal images of an olfactory bulb slice of GCaMP6f (Left), ChR2-mCherry (Center) and merged signals (Right) expression in DAT+ neurons. **(e)** Width of the exponential fits in (**Fig**. **4c**) against spots of same size and varying light intensity. Dots correspond to individual ROIs. We tested a range of photo-stimulation light intensities ranging from 0.5 to 10.0 mW/mm$^2$. Within the range of light intensity used, the characteristic width constant varied from 25 µm for the lowest to 62 µm for the highest.*

#### Supplementary Figure 7

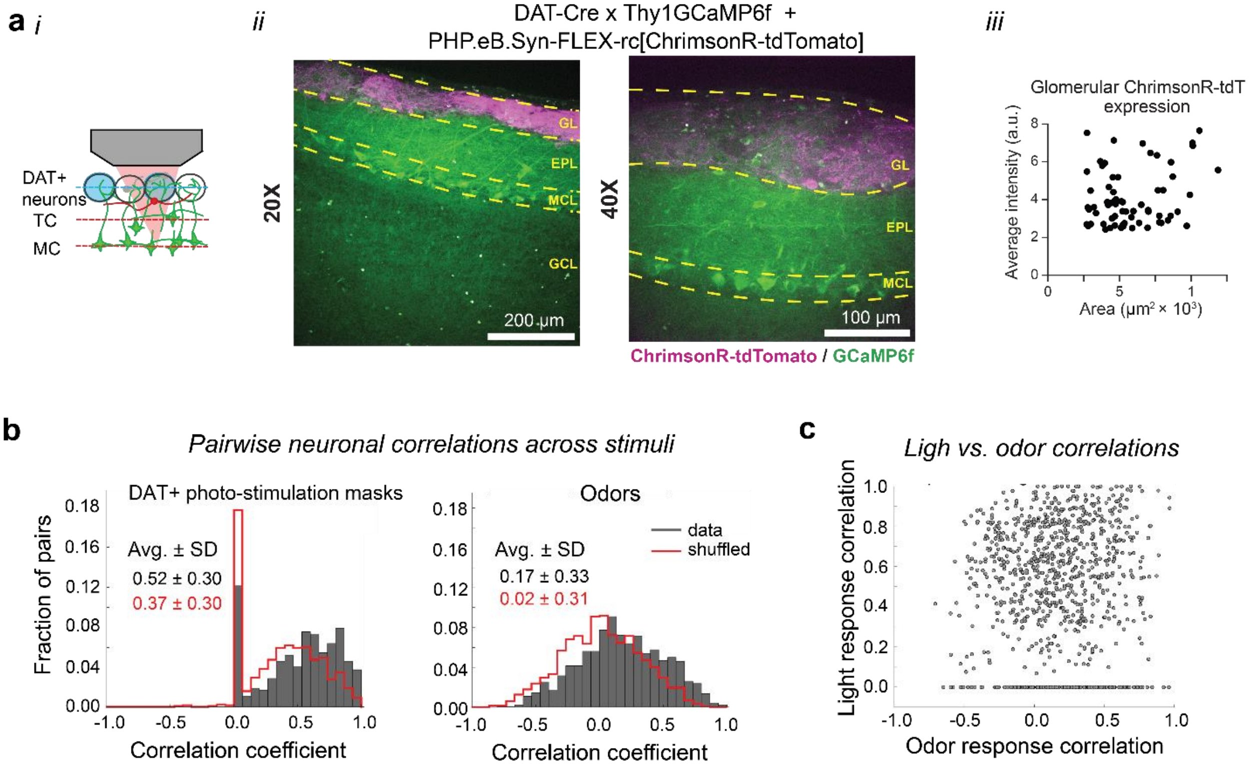


***Figure S7. Optogenetic control of DAT+ (superficial short axon) cells to determine their impact on mitral cell responses**. Corresponding to **Fig**. 4. **(a)** (i) Cartoon schematics: in vivo photo-stimulation of opsin expressing DAT+ cells while monitoring mitral cell activity (GCaMP6f) by axial decoupling of the photo-stimulation and imaging planes. (ii) Confocal image of olfactory bulb sagittal slices showing expression of ChrimsonR-tdtomato in the glomerular DAT+ cells (superficial short axon cells) and GCaMP6f in the mitral and tufted cells in DAT-Cre x Thy1GCaMP6f (5.11) mice. (iii) td-tomato intensity across glomeruli as a function of glomerular size (area). **(b)** Pairwise correlation of mitral cell responses across stimuli in an example field of view (correlation between all pairs of cells, corresponding to **Fig. 4h**). (Left) Example distribution of pairwise correlations of mitral cells responses across DAT+ cell glomerular photo-stimulation masks (G1-G6); data: 0.52 ± 0.30; shuffled: 0.37 ± 0.30. (Right) Example distribution of pairwise correlation of same cells (N= 49 mitral cells) in response to different odors in the panel (1-11). data: 0.17 ± 0.33; shuffled: 0.02 ± 0.31. Shuffled mitral cell index control distributions are shown in red. **(c)** Pairwise correlation of mitral cell responses to different DAT+ cell glomerular photo-stimulation masks (G1-G6) vs. pairwise correlation of same mitral cell responses to odors in the panel (1-11). Each dot corresponds to a pair of mitral cells. Responses across light stimuli appeared more similar than across odors at the level of mitral cell pairs. However, for this small panel of stimuli, no apparent trend emerged when analyzing the response correlation of a given pair of mitral cells responded to light stimulation of DAT+ glomerular masks as a function of their response similarity across the odors in the panel.*

#### Supplementary Figure 8

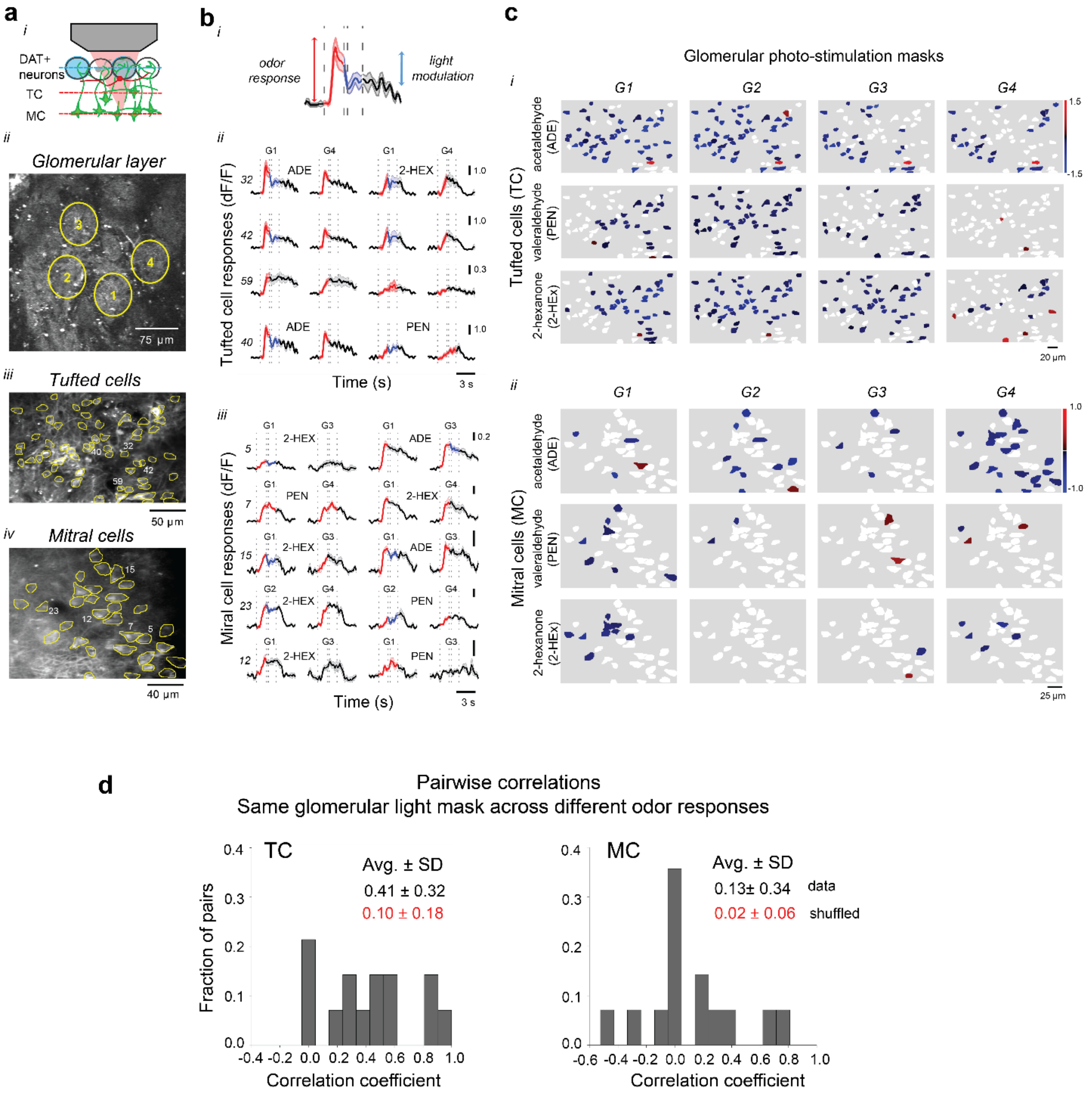


***Figure S8. Specificity of modulation of mitral and tufted odor responses by glomerular photo-activation of DAT+ interneurons.*** *Corresponding to* ***Fig****. 5.* ***(a)*** *(i) Schematic of the experiment: in vivo photo-stimulation of ChrimsonR-tdtomato expressing DAT+ cells, while monitoring its impact on mitral and tufted cell odor responses (GCaMP6f) using ADePT; (ii) baseline 2p fluorescence of an example field of stimulation in glomerular layer, and of two optical z-planes sampling tufted (iii) and mitral cells (iv) across different depths (100 µm vs. 275 µm from surface) in the same imaging session; circles and numbers indicate the location of projected light masks (ii) and example tufted and mitral cell bodies (iii,iv).* ***(b)*** *(i) Fluorescence change (ΔF/F) elicited*

*by odor presentation in one example tufted cell, and its subsequent modulation by projecting of a glomerular light mask to optogenetically boost DAT+ cell activity (500 ms, 10 mW/mm$^2$) 2s from odor onset. (ii, iii) Modulation of example tufted (ii) and mitral cell (iii) odor responses by photo-stimulating DAT+ interneurons across different glomeruli (G1-G4); (**c**) Odor response modulation (ΔF/F; two dimensional light suppression/activation maps) of tufted (i) and mitral cell (ii) odor responses (ADE, PEN, 2-HEX) evoked by glomerular photo-stimulation of DAT+ cells across 4 individual glomeruli (G1-G4); each cell is color coded (blue-to-red) to indicate its response amplitude to light/odor stimulation; non-significant responses were thresholded to white (Methods). (**d**) Example distribution of pairwise correlation of tufted (Left) and mitral (Right) cell response modulation for different odors across the same DAT+ cells glomerular photo-stimulation mask, aggregated across 4 masks (G1-G4; TC: 0.41 ± 0.32, Avg. ± SD; 14 pairs, 2 FOVs, 64 TCs; MC: 0.13±0.34; 14 pairs 2 FOVs, 39 MC cells). Avg. ± SD for shuffled cell index control distributions are shown in red.*
